# Deciphering the dynamical origin of mixed population during neural stem cell developmental lineage commitment

**DOI:** 10.1101/099903

**Authors:** Dola Sengupta, Sandip Kar

## Abstract

Neural stem cells (NSC's) often give rise to mixed population of cells during differentiation. However, the dynamical origin of these mixed states is poorly understood. In this article, our mathematical modeling study demonstrates that the bone morphogenetic protein 2 (BMP2) driven differential differentiation dynamics of NSC's in central and peripheral nervous systems essentially function through two distinct bi-stable switches that are mutually interconnected. Stochastic simulations of the model reveal that the mixed population originates due to the existence of these bistable switching regulations while the maintenance of such mixed states depends on the level of stochastic fluctuations of the system. Importantly, the model predicts that by individually altering the expression level of key regulatory proteins, the NSC's can be converted entirely to a preferred phenotype for BMP2 doses that earlier resulted into mixed population. Our findings show that efficient neuronal regeneration can be achieved by systematically maneuvering the differentiation dynamics.

**One-sentence summary:** Unraveling the differential dynamical origin, maintenance and escape route of mixed population in the midst of developmental fate commitment in central and peripheral nervous systems

## Introduction

Transformation of neural stem cells (NSC) exclusively to neuronal cells in a factor dependent manner is one of the most challenging aspects in modern era of stem cell research especially in the context of cell-based therapies. In this regard, Bone morphogenetic protein 2 (BMP2) can play a crucial role, as it is known to differentially regulate the differentiation of NSC's in central and peripheral nervous systems (*1–9*). In central nervous system (CNS), a low level of BMP2 drives the NSC's preferably towards neuronal fate (*1–3, 10–12*). Whereas, similar low dose of BMP2 produces glial cells from NSC's of peripheral nervous system (PNS) origin and only very high level of BMP2 can selectively effect neuronal transformation (*4–8*). To make the issue even more intriguing, for certain range of BMP2 doses, mixed populations of neuron and glial cells were obtained from cultured NSC's of both central and peripheral origins (*4–6*, *8*, *9*, *13–20*). What is the origin of these mixed states? How they are maintained in the cell culture medium in a BMP2 dose dependent manner? Furthermore, how to avoid such mixed population states for the same range of BMP2 doses to achieve unidirectional cell fate commitment? Even a qualitative answer to these questions can be extremely insightful to develop methods to expand NSC's specifically into preferred phenotypes.

BMP2 mediated lineage commitment of NSC's is an extremely complex event and involves a number of key transcription factors. Recent deterministic mathematical modeling study has revealed that the differential nature of the BMP2 driven differentiation in central and peripheral nervous systems is essentially governed by a mushroom kind of bifurcation observed in the steady state dynamics of these key cell fate regulatory transcription factors (like Mash1, mammalian achaete-scute homologue) as a function of increasing doses of BMP2 (*21*). More importantly, the mushroom kind of bifurcation feature does indicate that the intrinsic and extrinsic fluctuations present in and around the NSC's in the BMP2 culture medium may eventually lead to a mixed population of cells in a cell-type specific manner (*21*). Experiments performed with NSC's in BMP2 and other growth factor conditions do suggest that stochastic fluctuations may play a significant role to alter the cell fate decision-making process and can help to maintain a mixed population state (*4–6*, *8*, *9*, *14–19*, *22–24*). Thus, understanding the role of stochastic fluctuations in these systems may improve our understanding about the underlying regulatory dynamics of differentiation in the NSC's.

Mathematical and computational modeling was found to be quite useful and intuitive to analyze the effect of such kind of fluctuations qualitatively for biological systems of varied complexities (*25*, *26*). Keeping this in mind, we put forward a stochastic mathematical model of a minimal gene interaction network (consisting of Mash1, Id1 (inhibitor of differentiation) and E47 protein) that portrays the differential dynamics of differentiation of NSC's in CNS and PNS in a BMP2 dose dependent manner. We initially set up the stochastic model for the NSC's of CNS origin. By systematic sensitivity analysis of the model parameters and using bifurcation analysis, we further develop the stochastic model that describes the BMP2 driven dynamics of NSC's for a PNS like system. The stochastic analysis of the minimal model, where we have included both the intrinsic and extrinsic variability's, evidently shows how mixed population of neuron and glial cells can coexist at different extent under certain experimentally feasible range of BMP2 doses for the wild type (WT) NSC's for both CNS and PNS in a differential manner. Stochastic calculations demonstrate that the co-existence and extent of mixed population of cells will also depend on the ways the BMP2 level is altered (*suddenly* or *gradually*) during the experiment. The bifurcation analysis and the stochastic simulations of the minimal model further showed how by altering the expression level of E47 protein (*2*) or by lowering the expression level of Mash1 by over expression a particular microRNA (mir-34a) (*27*), the mixed population of cells could in principle be transformed to a preferred phenotype in a context dependent manner.

## Model construction

The minimal gene regulatory network, shown in Fig 1, describes how BMP2 controls the developmental fate in the NSC's of CNS and PNS by regulating the transcription of both Mash1 and Id1 proteins through direct and indirect pathways (*1*, *2*, *4*, *5*, *8*, *28*). Mash1 forms hetero-dimer with ubiquitously expressed bHLH protein E47 via their HLH domain (*1*, *2*, *10*). This heterodimer formation promotes CK2 mediated phosphorylation of Mash1 on Ser^152^ (*2*).

**Fig 1.**
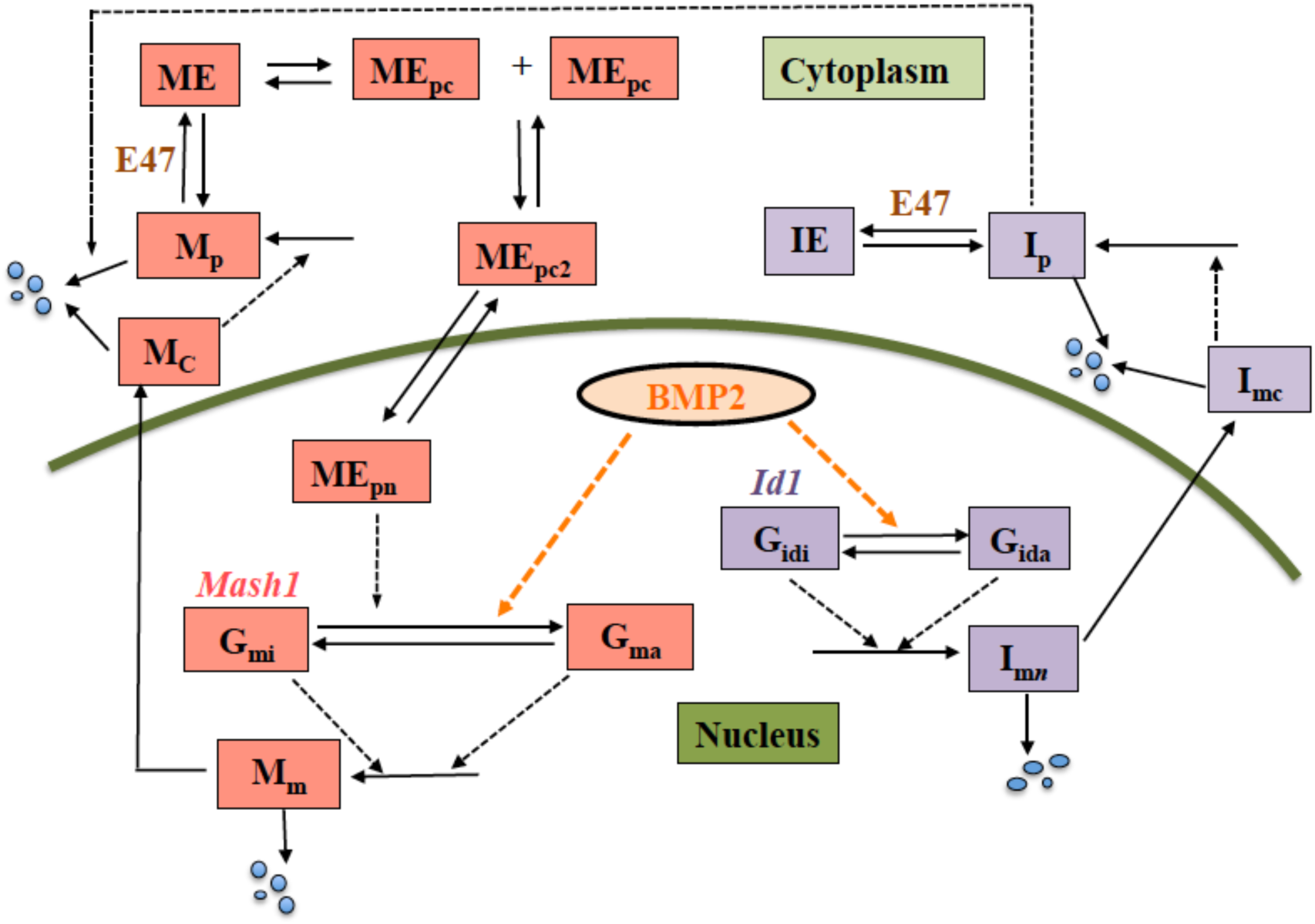
BMP2-driven neuronal differentiation regulation network. (Solid and dashed arrows represent direct and indirect activation processes respectively.) Once produced, *Mash1* mRNA (M_m_) gets transported from the nucleus to cytoplasm. Cytoplasmic *Mash1* mRNA (M_C_) produces Mash1 protein (M_p_) and it forms a heterodimer (ME) with E47 protein. After subsequent phosphorylation of Mash1 in ME, the phosphorylated component (ME_pc_) forms a dimer ME_pc2_. ME_pc2_ gets trans-located in to the nucleus and in the model it is designated as MEpn, which contributes to the *Mash1* gene activation in a positive feedback manner. Similarly *Id1* mRNA (I_mn_) is transported from nucleus to cytoplasm, where translation of Id1 protein (I_p_) takes place from its mRNA (I_mc_). I_p_ inhibits Mash1 through sequestration mechanism by forming an inactive complex (IE) with E47. I_p_ also promotes the degradation of M_p_. Corresponding kinetic equations, description of the variables and parameters are depicted in Table 1, SI Table and Table S2 respectively.

The phosphorylation of Mash1 in the hetero-dimeric complex leads to subsequent homo-dimerization of the hetero-dimeric complexes, which ultimately translocate back to the nucleus to bind to the promoter region of Mash1 gene (*21*). In this way Mash1 induces and maintains its own expression by reinforcing responsiveness to BMP2 via a positive-feedback loop (*1*, *2*, *4*, *10*). This BMP2 mediated positive feedback loop of Mash1 drives neuronal fate commitment in NSC's (*4*, *21*,*29*). On the other hand, BMP2 stimulation is also known to up regulate negative HLH-factor Id1 expression (*1*). Id1 antagonizes Mash1 in two ways. Firstly, Id1 sequesters E47 protein away from Mash1 and competitively binds with E47 protein that produces a transcriptionally inactive complex (*2*). Secondly, Id1 promotes degradation of the monomeric form of the Mash1 protein (*2*). Thus, Mash1 protein stability is critically regulated by E47/Id1 expression ratio (*2*).

The gene regulatory network depicted in Fig. 1 is translated into a deterministic mathematical model as given in Table 1 where all the differential equations (Table 1) for various species are solely based on mass action kinetics. The Variables are designated in Table S1. Description of the parameters, their numerical values and sources (*29*–*32*) are mentioned in Table S2.

**Table 1:**
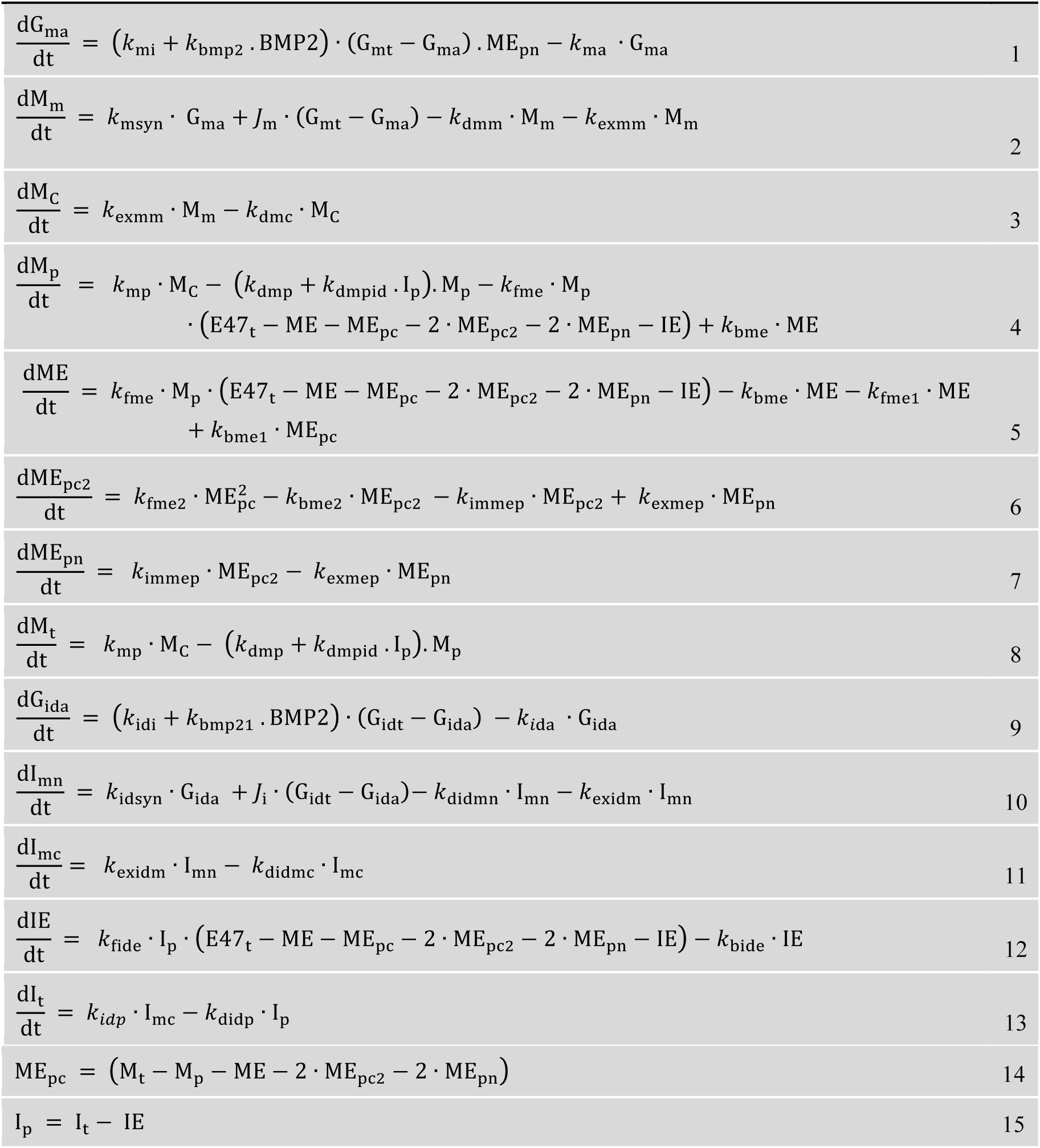
Equations governing the gene regulatory network of differentiation of NSC's.

|  |  |
| --- | --- |
| $\frac{dG_{ma}}{dt} = (k_{mi} + k_{bmp2} \cdot BMP2) \cdot (G_{mt} - G_{ma}) \cdot ME_{pn} - k_{ma} \cdot G_{ma}$ | 1 |
| $\frac{dM_m}{dt} = k_{msyn} \cdot G_{ma} + J_m \cdot (G_{mt} - G_{ma}) - k_{dmm} \cdot M_m - k_{exmm} \cdot M_m$ | 2 |
| $\frac{dM_c}{dt} = k_{exmm} \cdot M_m - k_{dmc} \cdot M_c$ | 3 |
| $\frac{dM_p}{dt} = k_{mp} \cdot M_c - (k_{dmp} + k_{dmpid} \cdot I_p) \cdot M_p - k_{fme} \cdot M_p \cdot (E47_t - ME - ME_{pc} - 2 \cdot ME_{pc2} - 2 \cdot ME_{pn} - IE) + k_{bme} \cdot ME$ | 4 |
| $\frac{dME}{dt} = k_{fme} \cdot M_p \cdot (E47_t - ME - ME_{pc} - 2 \cdot ME_{pc2} - 2 \cdot ME_{pn} - IE) - k_{bme} \cdot ME - k_{fme1} \cdot ME + k_{bme1} \cdot ME_{pc}$ | 5 |
| $\frac{dME_{pc2}}{dt} = k_{fme2} \cdot ME_{pc}^2 - k_{bme2} \cdot ME_{pc2} - k_{immep} \cdot ME_{pc2} + k_{exmep} \cdot ME_{pn}$ | 6 |
| $\frac{dME_{pn}}{dt} = k_{immep} \cdot ME_{pc2} - k_{exmep} \cdot ME_{pn}$ | 7 |
| $\frac{dM_t}{dt} = k_{mp} \cdot M_c - (k_{dmp} + k_{dmpid} \cdot I_p) \cdot M_p$ | 8 |
| $\frac{dG_{ida}}{dt} = (k_{idi} + k_{bmp21} \cdot BMP2) \cdot (G_{idt} - G_{ida}) - k_{ida} \cdot G_{ida}$ | 9 |
| $\frac{dI_{mn}}{dt} = k_{idsyn} \cdot G_{ida} + J_i \cdot (G_{idt} - G_{ida}) - k_{didmn} \cdot I_{mn} - k_{exidm} \cdot I_{mn}$ | 10 |
| $\frac{dI_{mc}}{dt} = k_{exidm} \cdot I_{mn} - k_{didmc} \cdot I_{mc}$ | 11 |
| $\frac{dIE}{dt} = k_{fide} \cdot I_p \cdot (E47_t - ME - ME_{pc} - 2 \cdot ME_{pc2} - 2 \cdot ME_{pn} - IE) - k_{bide} \cdot IE$ | 12 |
| $\frac{dI_t}{dt} = k_{idp} \cdot I_{mc} - k_{didp} \cdot I_p$ | 13 |
| $ME_{pc} = (M_t - M_p - ME - 2 \cdot ME_{pc2} - 2 \cdot ME_{pn})$ | 14 |
| $I_p = I_t - IE$ | 15 |

Since the deterministic model is constructed in terms of mass action kinetics, it can readily be used for stochastic calculation by employing Gillespie's stochastic algorithm (*33*) to understand the effect of intrinsic fluctuation present in the system. In our stochastic model, we added the effect of extrinsic fluctuations present due to difference in protein and mRNA numbers usually observed in an asynchronous population of cells in any culture medium (*34*–*36*). This kind of sets the platform to simulate the fluctuations present in the differentiation dynamics of the NSC's in a BMP2 culture medium and to analyze qualitatively the origin of the mixed populations of cells that is found in the experiments (*4*–*6*, *20*).

## Results

### Bmp2 driven stochastic developmental dynamics in WT NSC's of CNS origin

The bifurcation analysis of the model proposed in Table 1 reveals that the steady state levels of the total Mash1 protein concentration as a function of increasing BMP2 dose contains two interconnected bi-stable switches and gives rise to a mushroom kind of bifurcation (Fig 2a and Fig. S1). The saddle nodes SN_1_ and SN_2_ in this mushroom bifurcation (Fig 2a and Fig. S1) appear for very low doses of BMP2 (~ 0.041 s.u. and ~ 0.016 s.u.). Transcription of both *Mash1* and *Id1* genes are positively influenced by BMP2. Thus the appearance of a bi-stable switch for a very low level of BMP2 is possibly due to the fact that initially low levels of BMP2 induces E47 mediated *Mash1* transcription to a greater extent than the *Id1* transcription and causes the neuronal fate commitment (high level of Mash1) of NSC's at lower level of BMP2 in CNS. Subsequent increase in Bmp2 induces greater accumulation of Id1 protein that ultimately starts to affect the Mash1 expression level and consequently leads to the 2^nd^ bi-stable region with saddle nodes SN_3_ (BMP2 ~ 25.24 s.u.) and SN_4_ (BMP2 ~ 5.99 s.u.) (Fig. 2a and Fig. S1). Further increase in the BMP2 levels finally drives the system towards gliogenic fate.

**Fig 2.**
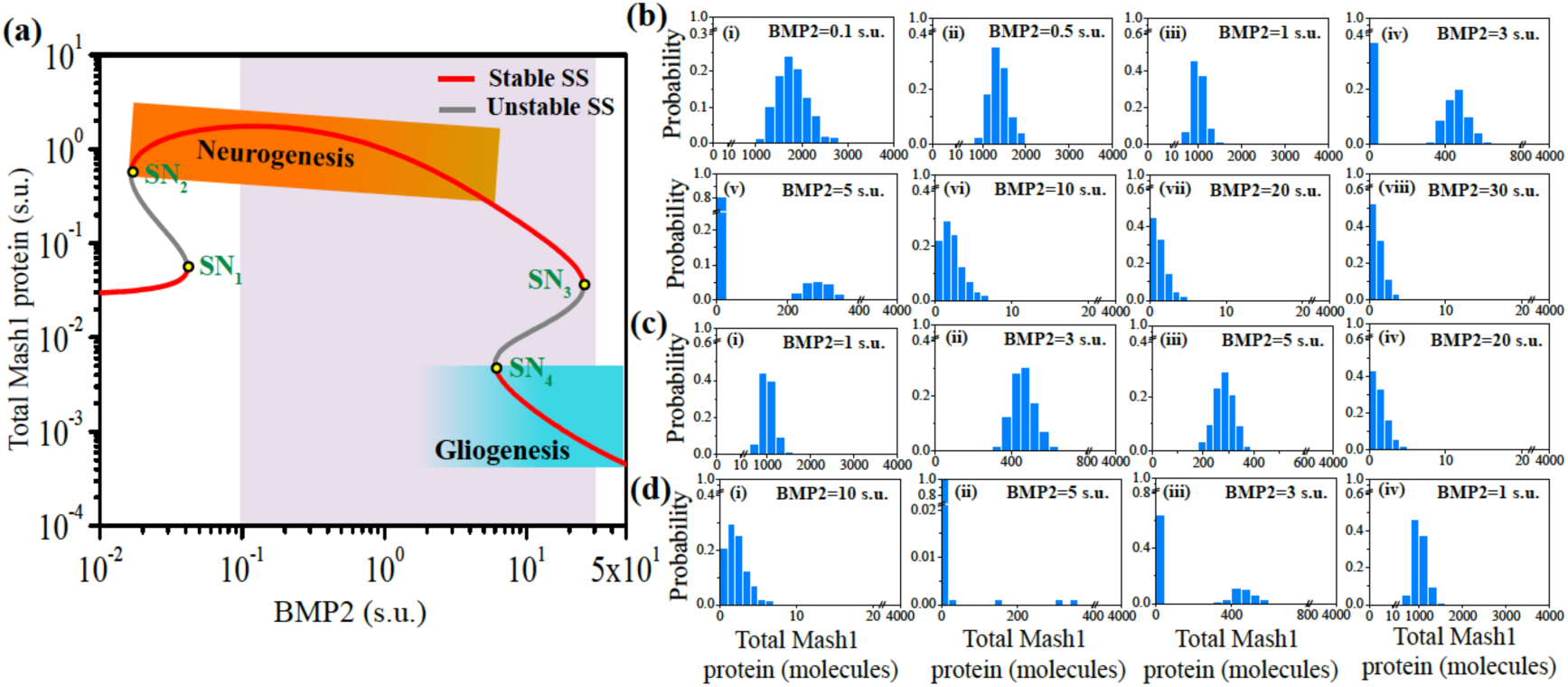
Bmp2 driven stochastic developmental dynamics in WT NSC's of CNS origin. **(a) Bifurcation diagram of total Mash1 protein is plotted as a function of BMP2** (both the axes are shown in log scale). Increasing level of BMP2 switches the developmental cell fate from neurogenic state (high Mash1 expression level, shown by the graded orange region) to gliogenic state (low Mash1 expression level, shown by the graded blue region). **Steady state distributions of total Mash1 protein are plotted at different doses of BMP2. (b)** BMP2 concentration in the cell culture medium is *suddenly increased* from 0.0005 s.u. to either **(i)** 0.1 s.u., or **(ii)** 0.5 s.u., or **(iii)** 1 s.u., or **(iv)** 3 s.u., or **(v)** 5 s.u., or **(vi)** 10 s.u., or **(vii)** 20 s.u., or **(viii)** 30 s.u.. **(c)** BMP2 concentration in the cell culture medium is *gradually increased* from 0.0005 s.u. to 0.1 s.u. and then from 0.1 s.u. to 1 s.u. **(i)** to **(ii)** 3 s.u., **(iii)** 5 s.u. and **(iv)** 20 s.u.. **(d)** BMP2 concentration in the cell culture medium is *gradually decreased* from 30 s.u. to 10 s.u. **(i)** to **(ii)** 5 s.u., **(iii)** 3 s.u. and **(iv**) 1 s.u.. All the simulations are done for 1000 cells and for four days (for details see methods section). To further confirm the results stochastic simulations are done for 2000 cells as well (Fig. S2).

At this point we performed the stochastic simulation of the model stated in Table 1 by implementing the Gillespie's stochastic simulation algorithm (SSA) (*33*) (using mass-action kinetic reactions provided in Table S3) for different concentrations of BMP2. We systematically incorporated the effect of extrinsic fluctuations by considering the fact that in an asynchronous population of cells the concentrations of different proteins and mRNA can vary from cell to cell (for details see method section) (*34*–*36*). While performing the stochastic simulations with different doses of BMP2, we stress on the fact that experimentally the BMP2 concentrations can be altered in two different ways that is either *suddenly* or *gradually.* We have considered both the situations one by one in our stochastic calculations.

In the first case (Fig. 2b), we have increased the BMP2 concentration in the cell culture medium *suddenly* to a higher value from the basal BMP2 concentration (0.0005 s.u.) needed to just maintain the cell culture. From the distribution plots (after 4 days of BMP2 stimulation) of the total Mash1 protein expression level obtained from the stochastic simulation results it is evident that mixed population of cells do exist when the BMP2 level is changed suddenly to 3 s.u. (Fig. 2b(iv)) and 5 s.u. (Fig. 2b(v)) from 0.0005 s.u.. On the other hand, *sudden* change of BMP2 concentration lower than 3 s.u. (either 0.1 s.u. (Fig. 2b(i)) or 0.5 s.u. (Fig. 2b(ii)) or 1 s.u. (Fig. 2b(iii))) leads to preferential neuronal fate commitment of NSC's in the cell culture medium. Whereas, BMP2 doses suddenly fixed higher than the 10 s.u. (either 10 s.u. (Fig. 2b(vi)) or 20 s.u. (Fig. 2b(vii)) or 30 s.u. (Fig. 2b(viii))) direct the NSC's in the cell culture medium completely towards gliogenic fate. Nakashima et al. had observed similar differentiation pattern when they increased the BMP2 concentration suddenly for the NSC's of CNS origin (*1*).

Instead of *sudden* increase, the experiment of rising the BMP2 dose in the cell culture medium can also be accomplished by *gradually* rising the BMP2 level in the same culture plate after every 4 days of interval. We performed an equivalent theoretical stochastic calculation where initially we have suddenly increased the BMP2 concentration from the basal level (0.0005 s.u.) to 0.1 s.u. (Fig. 2b(i)) to 1 s.u. (Fig. 2c(i)) and simulated the system for 96 hours. After that, we have *gradually* increased the BMP2 level from 1 s.u. to subsequently different higher levels (3 s.u. (Fig. 2c(ii)),5 s.u. (Fig. 2c(iii)) and 20 s.u (Fig. 2c(iv)) and simulated the system each time for 96 hours to get the distribution of total Mash1 expression level. Interestingly, for the intermediate BMP2 doses (i.e., 3 s.u. (Fig. 2c(ii)) and 5 s.u. (Fig. 2c(iii))) we did not get any mixed population (as observed in Fig. 2b(iv) and (v)) and the NSC's seem to choose the neuronal state with the higher Mash1 level. In an expected note, further gradual increase in the BMP2 concentration to 20 s.u. guides the NSC's towards gliogenic fate commitment. On the other hand, our stochastic simulations predict that if one decreases the BMP2 concentration gradually from 30 s.u. (Fig. 2b(viii)) to 10 s.u. (Fig. 2d(i)) to subsequent lower values of BMP2 (for example, 5 s.u. (Fig. 2d(ii)) and 3 s.u. (Fig. 2d(iii))) consecutively after each 96 hours of simulation, we will obtain bimodal Mash1 distributions for each concentration of BMP2 suggesting existence of mixed population under such circumstances. Further gradual decrease in the BMP2 concentration to 1 s.u. (Fig. 2d(iv)) from 3 s.u. steers the NSC's towards complete neuronal fate commitment.

These observations kind of highlight the fact that the way the experiments are performed in the laboratory may change the course of the ultimate cell fate decisions during BMP2 driven lineage commitment of NSC's in culture conditions and mixed population of cells can be avoided if we have better understanding about the underlying stochastic dynamics of the differentiation process. The question remains whether the NSC's of PNS origin shows similar BMP2 driven stochastic dynamics of differentiation or not?

### Bmp2 driven differential stochastic developmental dynamics in WT NSC's of PNS origin

To address the above-mentioned question, we first examine whether we can use the deterministic model proposed for NSC's of CNS origin (possibly with some minor adjustment) to describe the BMP2 driven differentiation dynamics of NSC's of PNS origin or not? We performed systematic sensitivity analysis of the model parameters by taking the position of the saddle nodes SN_1_ (Fig. S3) and SN_4_ (Fig. S4) separately as sensitivity criteria (For details see methods section). By analyzing the sensitivity plots we realized that there are BMP2-related pair of parameters (such as *k*_bmp2_ and *k*_bmp21_, Fig. S5a-b), which can be altered in a specific way to reconcile the BMP2 driven differentiation dynamics in NSC's of PNS like system. The biological relevance of choosing the parameters *k*_bmp2_ and *k*_bmp21_ has been discussed in SI Text (*1*, *6*, *37*–*39*).

The bifurcation analysis of the model given in Table 1 with changed values of *k*_bmp2_ and *k*_bmp21_ shows that for the NCS's of PNS origin the two interconnected bi-stable switches have significantly changed their relative positions (Fig. 3a and Fig. S5c) as compared to the CNS case (Fig. 2a). The saddle nodes SN_1_ and SN_2_ in this mushroom bifurcation (Fig. 3a and Fig. S5c) appear for relatively higher doses of BMP2 (~ 12.66 s.u. and ~ 3.94 s.u.) and the saddle nodes SN_3_ (BMP2 ~ 971.77 s.u.) and SN_4_ (BMP2 ~ 169.66 s.u.) are shifted to even higher levels of BMP2. Now if we consider the experimental range of BMP2 (as taken in case of CNS) between 0.1 s.u. (low value) to 30 s.u. (reasonably high value) then in that range the first bi-stable switch of the mushroom bifurcation will govern the cell fate decisions of the NSC's in a PNS like system.

**Fig 3.**
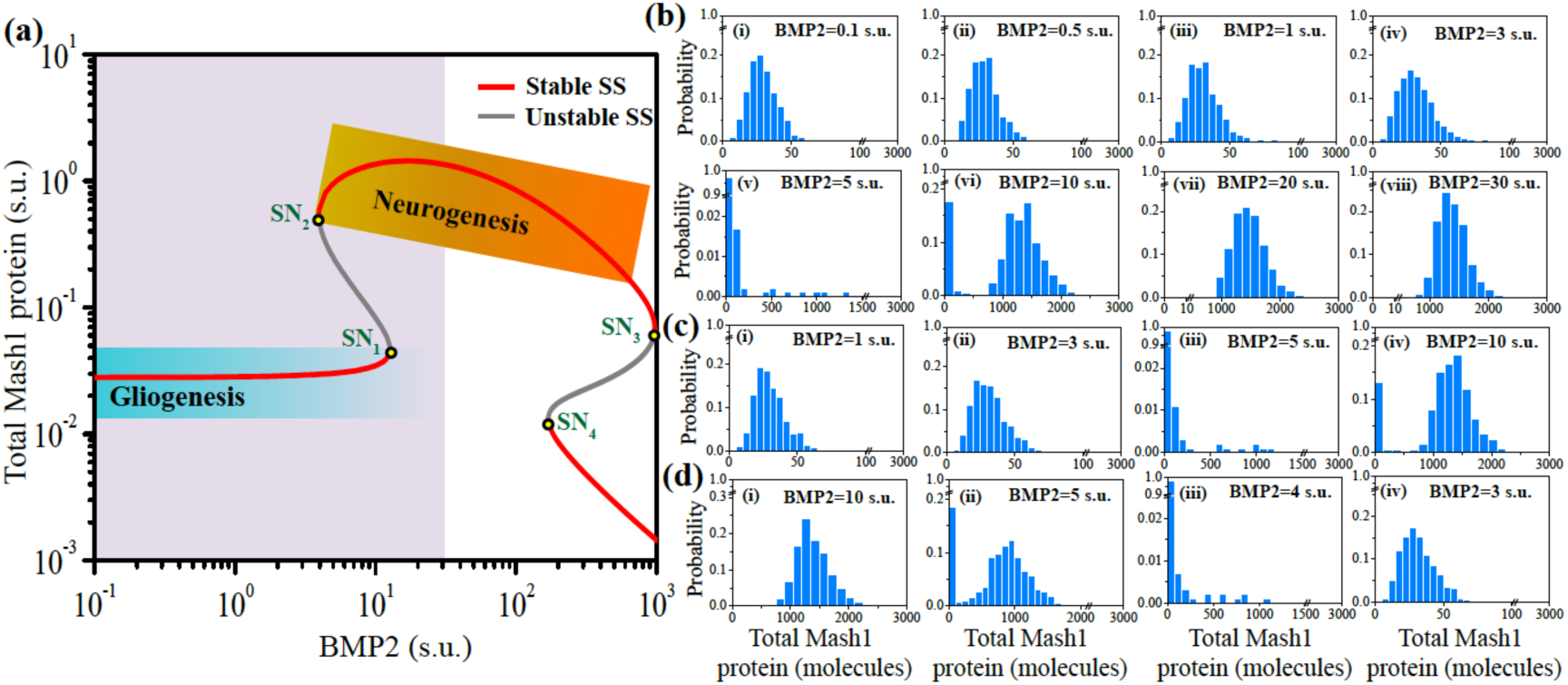
Bmp2 driven stochastic developmental dynamics in WT NSC's of PNS origin. **(a) Bifurcation diagram of total Mash1 protein is plotted as a function of BMP2** (both the axes are shown in log scale). Increasing level of BMP2 switches the developmental cell fate from gliogenic state (low Mash1 expression level, shown by the graded blue region) to neurogenic state (high Mash1 expression level, shown by the graded orange region). *k*_bmp2_=400 min^−1^ and *k*_bmp21_=1.2X10^−2^ min^−1^ are used to get PNS like feature. Other parameters are same as depicted in Table S2. **Steady state distributions of total Mash1 protein are plotted at different doses of BMP2. (b)** BMP2 concentration in the cell culture medium is ***suddenly increased*** from 0.0005 s.u. to either **(i)** 0.1 s.u., or **(ii)** 0.5 s.u., or **(iii)** 1 s.u., or **(iv)** 3 s.u., or **(v)** 5 s.u., or **(vi)** 10 s.u., or **(vii)** 20 s.u., or **(viii)** 30 s.u.. **(c)** BMP2 concentration in the cell culture medium is ***gradually increased*** from 0.0005 s.u. to 0.1 s.u. and then from 0.1 s.u. to 1 s.u. **(i)** to **(ii)** 3 s.u., **(iii)** 5 s.u. and **(iv)** 10 s.u.. **(d)** BMP2 concentration in the cell culture medium is ***gradually decreased*** from 30 s.u. to 10 s.u. **(i)** to **(ii)** 5 s.u., **(iii)** 4 s.u. and **(iv)** 3 s.u.. All the simulations are done for 1000 cells and for four days to corroborate with the experiments (for details see methods section). To further confirm the results stochastic simulations are done for 2000 cells as well (Fig. S6).

We perform the stochastic simulation by *suddenly* increasing BMP2 concentration (from basal level 0.0005 s.u.) in the cell culture medium to higher values and plotted the distribution of total Mash1 protein level (Fig. 3b(i-viii)) in each case after 4 days of BMP2 stimulation. We clearly observed mixed population of cells when the BMP2 level was changed suddenly to 10 s.u. (Fig. 3b(vi)). Partial mixed population was even spotted for *sudden* increase of BMP2 to 5 s.u. (Fig. 3b(v)). For BMP2 concentrations *Suddenly* fixed lower than 5 s.u. (either 0.1 s.u. (Fig. 3b(i)) or 0.5 s.u. (Fig. 3b(ii)) or 1 s.u. (Fig. 3b(iii)) or 3 s.u. (Fig. 3b(iv))), the NSC's of PNS origin in the cell culture medium exclusively opt for gliogenic fate. Whereas a *sudden* increase in the BMP2 dose higher than 10 s.u. (either 20 s.u. (Fig. 3b(vii)) or 30 s.u. (Fig.3b(viii))) favorably transform the NSC's to neuronal cells. Lo et al. had found similar fate commitment pattern and mixed population of neuronal and non-neuronal cells when they increased the BMP2 concentration suddenly for the NSC's of PNS origin in a systematic manner (*4*).

We further executed stochastic simulations where BMP2 concentration is elevated *gradually* from 0.1 s.u. (Fig. 3b(i)) to 1 s.u. (Fig. 3c(i)) and then from 1 s.u. (Fig. 3c(i)) to successively different higher levels (3 s.u. (Fig. 3c(ii)), 5 s.u. (Fig. 3c(iii)) and l0 s.u (Fig. 3c(iv)) by following the same method discussed earlier for CNS case. The simulation results confirm that we will get mixed population of cells for the BMP2 doses 5 s.u. and 10 s.u. in a similar manner as is found when the BMP2 concentration is changed *suddenly*. Intriguingly, if we start from a steady state distribution of cells for example after 96 hours of sudden BMP2 stimulation from 0.0005 s.u. to 30 s.u. (Fig. 3b(viii)) and gradually decrease the BMP2 level to 10 s.u. (Fig. 3d(i)), we do not observe a bimodal Mash1 distribution indicating that the cells remain solely committed to neuronal fate. Further *gradual* decrease in the BMP2 level to 5 s.u. (Fig. 3d(ii)) and 4 s.u. (Fig. 3d(iii)), reestablish the mixed population of cells in the cell culture medium that ultimately commits to only gliogenic fate once BMP2 level is reduced to 3 s.u. (Fig. 3d(iv)). These observations kind of reiterate the fact that the manner in which the BMP2 stimulation experiment is performed can influence the overall cell fate decision-making process in a culture medium even in case of a PNS like system as well.

### Model predicts the possible routes of unidirectional lineage commitment of NSC's by altering E47 expression level

Can the proposed model provide further insight by predicting the possible routes to drive the NSC's towards unidirectional lineage commitment for BMP2 doses where in WT situation we get mixed population? To investigate this issue in silico, we systematically performed over expression and knock down studies of the E47 and Id1 proteins respectively. Deterministic bifurcation analysis (Fig. 4a) suggests that by either over expressing (Fig. 4a, **1**) or knocking down (for example, Fig. 4a, **3**) the E47 protein, it is possible to transform the NSC's of CNS origin exclusively to either neuronal or glial cells under the same experimental doses of BMP2 where we have obtained mixed population of cells in WT situation (Fig. 4a, WT). Interestingly, the steady state profile of Mash1 protein as a function of BMP2 transits to an isola kind of bifurcation state (*40*) (Fig. 4a, **2** and **3**) from a mushroom like bifurcation (Fig. 4a, WT) as the E47 protein is down regulated in comparison to the WT situation. The bifurcation appears as the saddle nodes SN_1_ and SN_4_ eventually coalesce and annihilate (Fig. 4a, **2** and **3**) each other with a decreasing level of E47 protein. Whereas, an increase in the E47 protein level leads to the greater separation of the saddle nodes SN_1_ and SN_4_ (Fig. 4a, **1**) that increases the chance of preferential neurogenesis over mixed population (Fig. 2b(v)) say for 5 s.u. of BMP2 dose.

**Fig 4.**
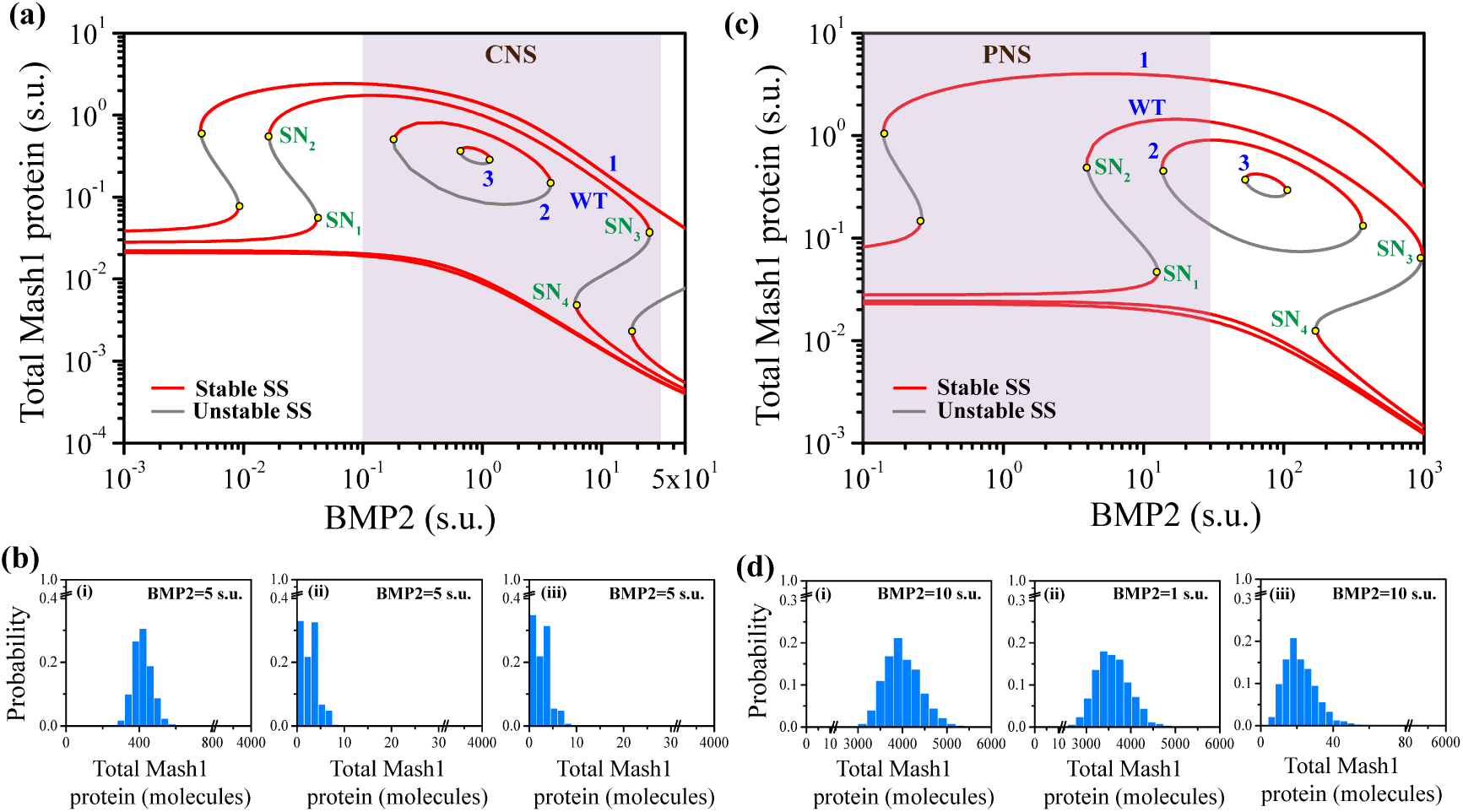
Alteration in E47 expression level could lead to unidirectional lineage commitment of NSC's in CNS and PNS. **(a)** Changes in the bifurcation of the total Mash1 protein from mushroom bifurcation (situation **WT**, E47_t_=0.6 s.u. for NSC's of CNS origin) either to (**1**) extended mushroom bifurcation (by increasing the E47_t_ level to 1.2 s.u.)), or to (**2** and **3**) an ‘isola’ kind of bifurcation (by decreasing the E47_t_ level to 0.2 s.u. and to 0.15 s.u. consecutively). **(b)** Distribution of total Mash1 protein concentration when BMP2 concentration in the cell culture medium is ***suddenly increased*** from 0.0005 s.u. to either (i) 5.0 s.u. (with E47_t_=1.2 s.u.) or, (ii) 5.0 s.u. (with E47_t_=0.2 s.u.) or, (iii) 5.0 s.u. (with E47_t_=0.15 s.u.). **(c)** Changes in the bifurcation of the total Mash1 protein from mushroom bifurcation (situation **WT**, E47_t_=0.6 s.u. for NSC's of PNS origin) either to (**1**) extended mushroom bifurcation (by increasing the E47_t_ level to 3.0 s.u.)), or to (**2** and **3**) an ‘isola’ kind of bifurcation (by decreasing the E47_t_ level to 0.35 s.u. and to 0.25 s.u. consecutively. **(d)** Distribution of total Mash1 protein concentration when BMP2 concentration in the cell culture medium is ***suddenly increased*** from 0.0005 s.u. to either (i) 10.0 s.u. (with E47_t_=3.0 s.u.) or (ii) 1.0 s.u. (with E47_t_=3.0 s.u.) or (iii) 10.0 s.u. (with E47_t_=0.25 s.u.). Other parameters are same as depicted in Table S2 and all the simulations are performed for 1000 cells (for details see method section).

Stochastic simulation of the model with over expressed E47 level for 5 s.u. of BMP2 dose (Fig. 4b(i)) conforms the deterministic bifurcation analysis as the NSC's of CNS origin preferentially opt for neuronal state under such condition, which is evident from a uni-modal distribution of Mash1 protein centered around high levels of Mash1. On the other hand, stochastic calculation for the same BMP2 dose (5 s.u.) with reduced level of E47 (E47_t_=0.2 s.u., Fig. 4b(ii), and E47_t_=0.15 s.u., Fig. 4b(iii)) confirms the exclusive glial fate commitment of NSC's with uni-modal distribution of Mash1 centered on low levels of Mash1. Intriguingly, stochastic simulations further indicate if the BMP2 level is suddenly increased to either 0.2 or 0.5 s.u. from basal level and after 96 hours the E47 is knocked down in the cell culture medium by some experimental means then there lies a possibility of observing a mixed population (Fig. S7a(i)-(ii)) even for an isola kind of situation (Fig. 4a, **2**). Although for a relatively higher level of BMP2 (2 s.u.) the NSC's will ultimately choose the glial state (Fig. S7a(iii)).

We did the bifurcation studies for the E47 over expression and knock down situations in a similar manner for the PNS like system. Here also we found that the E47 over expression could lead to unidirectional differentiation of NSC's towards Neuron (Fig. 4c, **1**) while the E47 knock down might favor the glial fate commitment (Fig. 4c, **2** and **3**). Stochastic calculations implemented for E47 over expressed situation (Fig. 4c, **1**) at 10 s.u. of BMP2 dose leads to unidirectional neuronal fate commitment (Fig. 4d(i)) instead of the mixed population as observed for the WT NSC's of PNS origin. The unidirectional fate commitment to neurons remains true (Fig. 4d(ii)) for even lower doses of BMP2 (1 s.u.) for the E47 over expressed case. Furthermore, the stochastic results obtained for E47 knock down situation for 10 s.u. of BMP2 gives rise to unidirectional fate commitment to glial cells (Fig. 4d(iii)) and corroborate nicely to the predictions made from the deterministic bifurcation study.

Moreover, the deterministic and stochastic simulations indicate that by either knocking out or over expressing *Id1* gene as performed in the experiments (*1*, *2*, *41–43*), one can preferentially get either neuron (Fig. S7b, for BMP2 = 10 s.u.) or glial (Fig. S7c, for BMP2 = 1 s.u.) cells in the BMP2 cell culture medium where in case of WT NSC's we have obtained exactly opposite cell types for the same BMP2 doses (Fig. 2b(iii) and Fig. 2b(vi)). In a similar fashion if the synthesis rate of Mash1 protein is reduced (say by over expressing microRNA's such as by Mir-34a (*27*)) or Id1 protein synthesis rate is increased numerically, the NSC's of CNS origin can be transformed to a single gliogenic state (Fig. S8) for the same BMP2 dose (say 5 s.u.) where mixed population state was observed in WT situation (Fig. 2b(v)). Thus, the modeling study reveals that cell fate decision making process in these NSC's of different origins can be nicely altered by influencing the steady state dynamics of the Mash1 protein level as a function of BMP2 dose.

## Discussion

BMP2 driven differentiation of NSC's specifically to either Neuron or Glial cells is an important and challenging issue to resolve as a better understanding to this can have definite therapeutic applications. Unfortunately, for a wide range of BMP2 doses, the NSC's maintain a mixed state with both neuronal and glial cells present at different extent in a culture medium and to make the matter even worse it happens in a differential manner for NSC's of CNS and PNS origin (*1–9*, *20*). In this article, by using a stochastic mathematical model (Table 1 and Table S3) of a minimal BMP2 driven differentiation regulatory network (Fig. 1), we intend to unravel the possible reasons behind the origin and maintenance of such mixed populations with an ultimate goal to foretell the probable avenues for obtaining specifically a particular differentiation phenotype by circumventing such mixed populations.

Our model qualitatively provided us with some crucial insights about the BMP2 driven differentiation dynamics that had been summarized schematically in Fig. 5. To begin with, bifurcation analysis of the model reveals that a mushroom kind of bifurcation observed in the Mash1 protein dynamics as a function of increasing doses of BMP2 controls the differentiation of the NSC's in CNS (Fig. 5a, left panel) and PNS (Fig. 5a, right panel). It further indicates that the differential nature of the differentiation of NSC's within a particular defined range of low (0.1 s.u.) and high (30 s.u.) dose of BMP2 can be explained if the overall differentiation dynamics is governed by the second (Fig. 2a) and the first (Fig. 3a) bi-stable switches respectively for CNS and PNS like system. The existence of mixed population for both CNS and PNS stems from the fact that a bi-stable switch in either of the situations governs the differentiation dynamics while the range of BMP2 for which the mixed population will be maintained is determined by the relative separation of the two consecutive saddle nodes (SN_3_ and SN_4_ for CNS and SN_1_ and SN_2_ for PNS). Our stochastic simulations that account for the intrinsic and extrinsic fluctuations present in the proposed differentiation network, qualitatively verified the presence of mixed population states (Fig. 2b-d, CNS and Fig. 3b-d, PNS) in the cell culture containing NSC's for a range of BMP2 doses.

**Fig 5.**
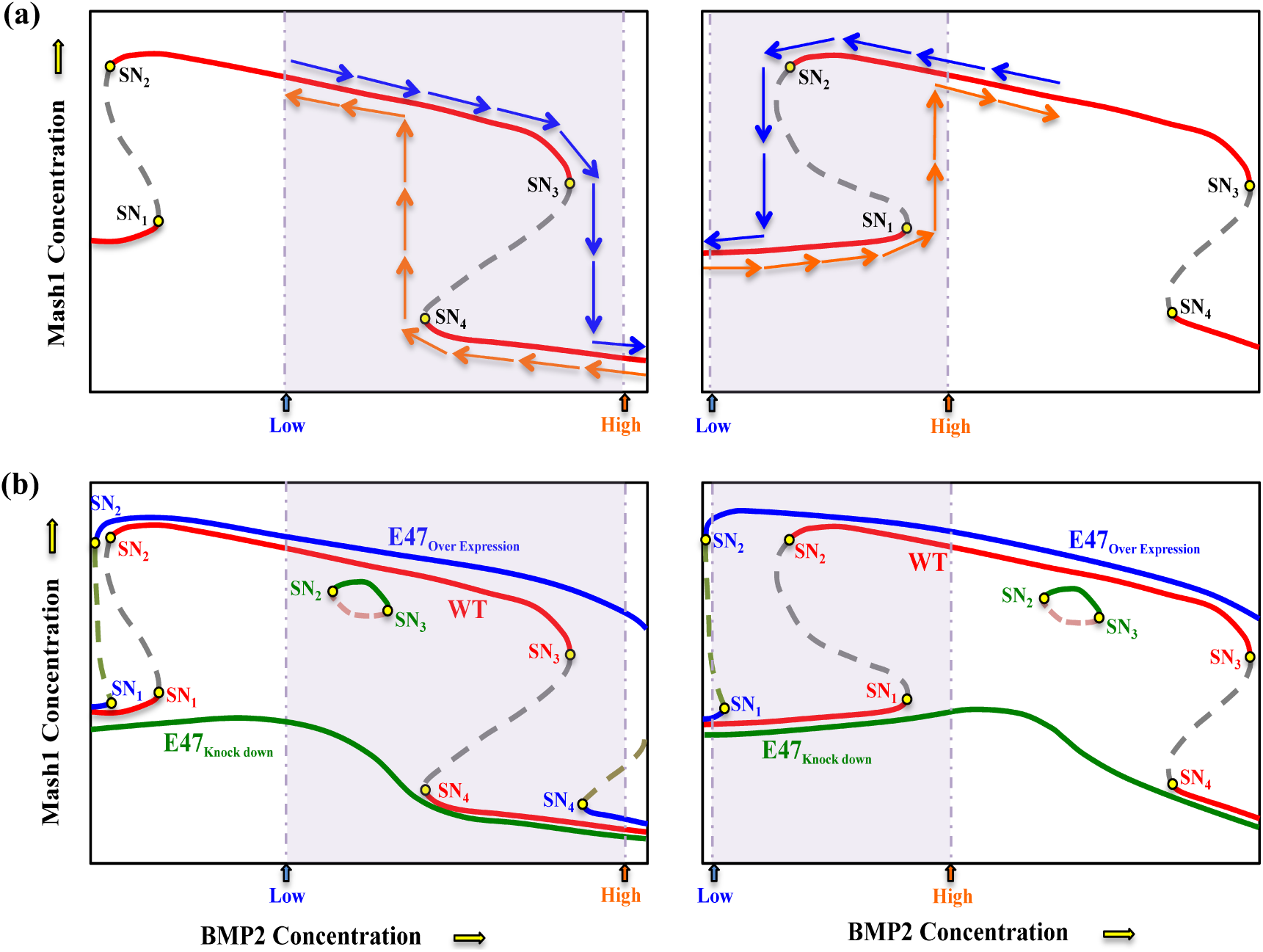
Model predicts about the origin, maintenance and possible escape routes from mixed population during BMP2 mediated differentiation. **(a)**The Mushroom bifurcation of Mash1 protein as a function of BMP2 governs the differential differentiation of NSC's. The 2^nd^ bi-stable domain (left panel) in Mushroom bifurcation leads to the mixed population of cells in case NSC's of CNS whereas the 1^st^ bi-stable switch (right panel) does the same for a PNS kind of system for a certain experimental range of BMP2 doses. The maintenance and extent of mixed states in the bistable range will depend on the fluctuations and relative separations of the steady states for both the cases. **(b)** One can evade the mixed state by either over expressing or down regulating the E47 protein level to direct the differentiation of NSC's to a specific phenotype in both CNS (left panel) and PNS (right panel) for a particular BMP2 dose.

Interestingly, the stochastic results suggest that the ratio of the individual cell types obtained within a mixed population state can vary depending on the way (i.e., *Suddenly* or *gradually)* BMP2 concentration is modified during experiment. It might happen that one can observe mixed state if the experiment is performed with *sudden* increase of BMP2 (Fig. 2b(v)) but a *gradual* increase at the same BMP2 level leads to a unidirectional lineage commitment (Fig. 2c(iii)). What are the possible reasons behind such observation? When BMP2 is suddenly increased to 5 s.u. from a very low dose (say 0.0005 s.u.), due to both intrinsic and extrinsic fluctuations present in the asynchronous cellular population in the cell culture medium we will initially obtain a mixed state. This bimodal distribution will eventually evolve with time and finally settle down to a particular distribution (for example, Fig. 2b(vi)) depending on the relative separation of the two stable steady states (shown in Fig. 2a) and the amount of fluctuation present in the system. On the other hand, for 1 s.u. of BMP2 the system was already preparing to be in a neural state with high Mash1 level (Fig. 2c(i)) and as BMP2 level is gradually increased to 5 s.u. the effect of fluctuation was not enough to take the cells to gliogenic state (i.e., in a low Mash1 state) resulting in a situation observed in Fig. 2c(iii). The role of fluctuations becomes even more evident when the in silico thought experiment with BMP2 *gradually* increased from 1 s.u. to 10 s.u. (Fig. 3c(iv)) and BMP2 *gradually* decreased to 10 s.u. (Fig. 3d(i)) from 30 s.u. gave rise to different cellular distributions for NSC's of PNS origin in a cell culture medium.

Our model can further predict the possible means to influence the dynamics of the BMP2 driven differentiation of NSC's by over expressing or knocking down the E47 protein (*2*) (Fig. 4). The schematic diagrams in Fig. 5b clearly demonstrate that one can avoid the existing bi-stable domain in the WT scenario for the NSC's in CNS (Fig. 5b, left panel) as well as in PNS (Fig. 5b, right panel) cases by either over expressing or knocking down E47 protein adequately in the experimentally employed range of BMP2 doses. On one hand, separation of the saddle nodes SN_1_ and SN_4_ considerably in the Mash1 steady state dynamics (Fig. 5b, left panel or right panel) controls the E47 over expression driven neuronal fate commitment, while annihilation of the saddle nodes (SN_1_ and SN_4_) with the decreasing levels of E47 through an isola kind of bifurcation (*40*) ensures the gliogenic state transformation. Similar kind of observations has been made for NSC's of CNS origin when Mash1 protein level is reduced (say by increasing Mir-34a expression (*27*)) or Id1 level is increased in our modeling studies (Fig. S8).

One can obviously ask whether we have enough experimental evidence in literature to support our qualitative modeling results or not? Our stochastic simulation results for NSC's of PNS origin (Fig. 3) do qualitatively corroborate with the experimental observations made by Lo et al. where they had shown that (i) mixed population of neural and glial cells do exist for a range of BMP2 doses and (ii) the extent of neuronal cells in the mixed population for different doses of BMP2 correlates with Mash1 concentration (*4*). Our modeling results (Fig. 2) additionally substantiate the differential nature of differentiation of the NSC's of CNS origin in the same experimental range of BMP2 doses as shown by Nakashima et al. (*1*). Unfortunately, they had not executed any experiment where they had increased the BMP2 concentration gradually in the cell culture medium to verify our bi-stable switch hypothesis controlling the fate commitment. Furthermore, there were evidences in literature (*2*) where E47 over expression had caused higher levels of Mash1 in NSC's of CNS origin and as a consequence neuronal state is favored over gliogenesis as qualitatively predicted by our deterministic and stochastic calculations (Fig. 4a-b). Not only that, our modeling results (Fig. S8) further corroborate the experimental observation that decreasing the Mash1 level by over expressing Mir-34a (*27*) will eventually cause a gliogenic fate commitment.

Our minimal stochastic model qualitatively provides valuable insight about the underlying fluctuating differentiation dynamics of NSC's with many novel predictions that remain to be validated experimentally. We are well aware of the fact that to make these calculations further quantitative, one requires more precise knowledge about the abundance levels of different proteins and mRNAs related to the network, which is currently not available for these systems. Moreover, implementing the concept of cell division during differentiation in the model may foster complete inclusion of the extrinsic fluctuation present in the system. Nonetheless, our model, even in its simple form, still nicely depicts the essential features of the BMP2 driven differential and stochastic dynamics of differentiation of the NSC's in CNS and PNS. It not only unravels how the mixed population of cells are originated and maintained for a range of BMP2 doses in the cell culture medium but also shows the avenues to elude the mixed population state to achieve unidirectional cell fate commitment. We believe that the lessons learned from this model will initiate novel experiments in these directions to advance our understanding about efficient neuronal regeneration.

## Materials and Methods

### Deterministic analysis

The mass-action kinetics based complete gene regulatory network (Fig. 1) was constructed in terms of 13 ordinary differential equations. The deterministic bifurcation analysis of the model was performed using the freely available software XPP-AUT. The bifurcation diagrams and time profiles were drawn with the help of OriginLab using the data points generated by XPP-AUT.

For the systematic sensitivity analysis, each parameter was increased individually at an amount of 20% with respect to the values given in Table S2 keeping all other parameters constant (Fig. S3 and Fig. S4). Purple colored bar signifies movement of the saddle nodes (SN_1_ and SN_4_) towards lower BMP2 level and light-grey colored bar signifies movement of the saddle nodes (SN_1_ and SN_4_) towards higher BMP2 than the WT CNS case.

### Stochastic simulations

The deterministic mathematical model, developed in terms of mass action kinetics, is readily translated into stochastic model by using the Gillespie's stochastic simulation algorithm (SSA) (*33*) that can capture the molecular fluctuation in a network quite adequately. For the SSA implementation (*33*), the propensity function of individual reactions of the deterministic mathematical model were accurately written in terms of number of molecules by introducing an appropriate conversion factor. The reactions and the propensities of the reactions are given in Table S3. In our stochastic simulation we wanted to take an ensemble average, for example, of a particular mRNA level present in each individual cell in an asynchronous cell culture medium.

To perform this, we assume that each individual SSA run with a particular starting seed of the uniformly distributed random number generator, corresponds to the trajectories of different proteins and mRNAs for an individual cell. In this way, we executed SSA simulation for 1000 individual cells. In addition, we initialized the dynamical system at very low dose of BMP2 (representative of cells in culture medium with almost no external BMP2 (BMP2=0.0005 s.u.)) and used those initial states as the initial condition in the each stochastic simulation run. The initial conditions were different for each run to account for the “extrinsic” noise due to cell-to-cell variability's (about 10 – 60% coefficient of variation (CV)) observed experimentally in an asynchronous population of cells for either mRNAs or proteins (34). We assumed that each mRNAs and proteins present in the network are distributed log-normally with the deterministic initial condition at BMP2=0.0005 s.u. as the mean and with about 20% CV (CV = 0.2) (*34–36*). Random initial conditions were then drawn from those log-normal molecular distributions of the individual variables with different initial means and 20% CVs before the start of the stochastic simulation for an individual cell. In this way the stochastic simulations were performed for individual cell traces having different concentrations of the network components that can account for the extrinsic noise along with the intrinsic noise present in the biochemical reactions (*34–36*). Stochastic simulations were done for 1000 cells (Figs 2, 3, 4 and Figs. S7 and S8) and 2000 cells (Figs. S2 and S6) and results seems to quite consistent so we stick to simulate for 1000 cells as it takes less computational time with no changes in the conclusions.

## Acknowledgements

Thanks are due to IRCC, IIT Bombay (13IRTAPSG005) for a fellowship to (DS). This work is supported by the funding agencies IRCC, IIT Bombay (13IRCCSG008), DST grant (EMR/2014/000500) and DBT grant (BT/PR11932/BRB/10/1315/2014).

## Supplementary Figures

**Fig. S1.**
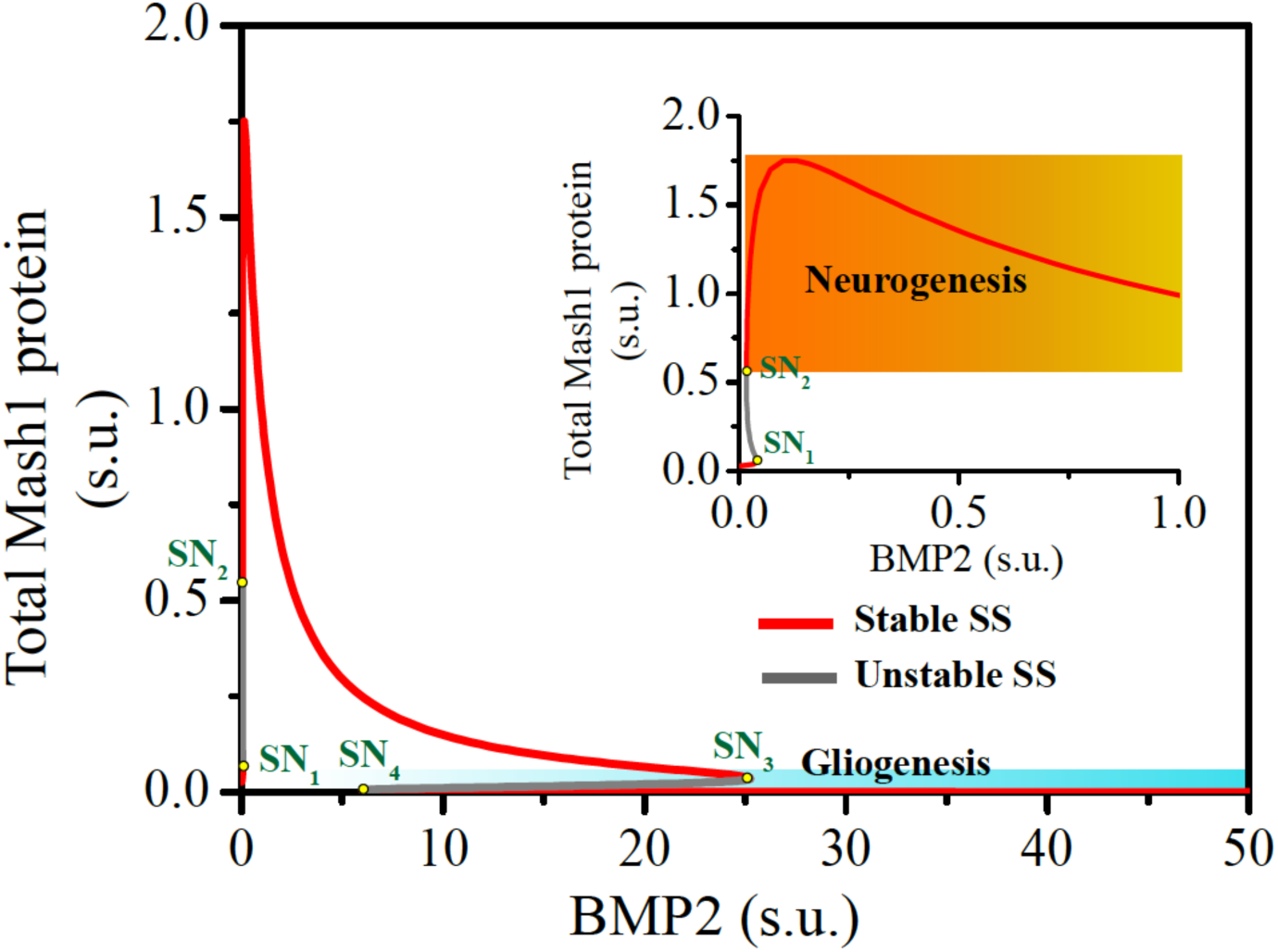
Bifurcation diagram of the total Mash1 protein steady state level as a function of BMP2 dose in CNS. Total Mash1 protein steady state is plotted as a function of BMP2 (both the axes are shown in linear scale). Increasing level of BMP2 switches the developmental cell fate from neurogenic state (high Mash1 expression level, shown by the graded orange region (inset)) to gliogenic state (low Mash1 expression level, shown by the graded blue region). Parameters are depicted in Table S2.

**Fig. S2.**
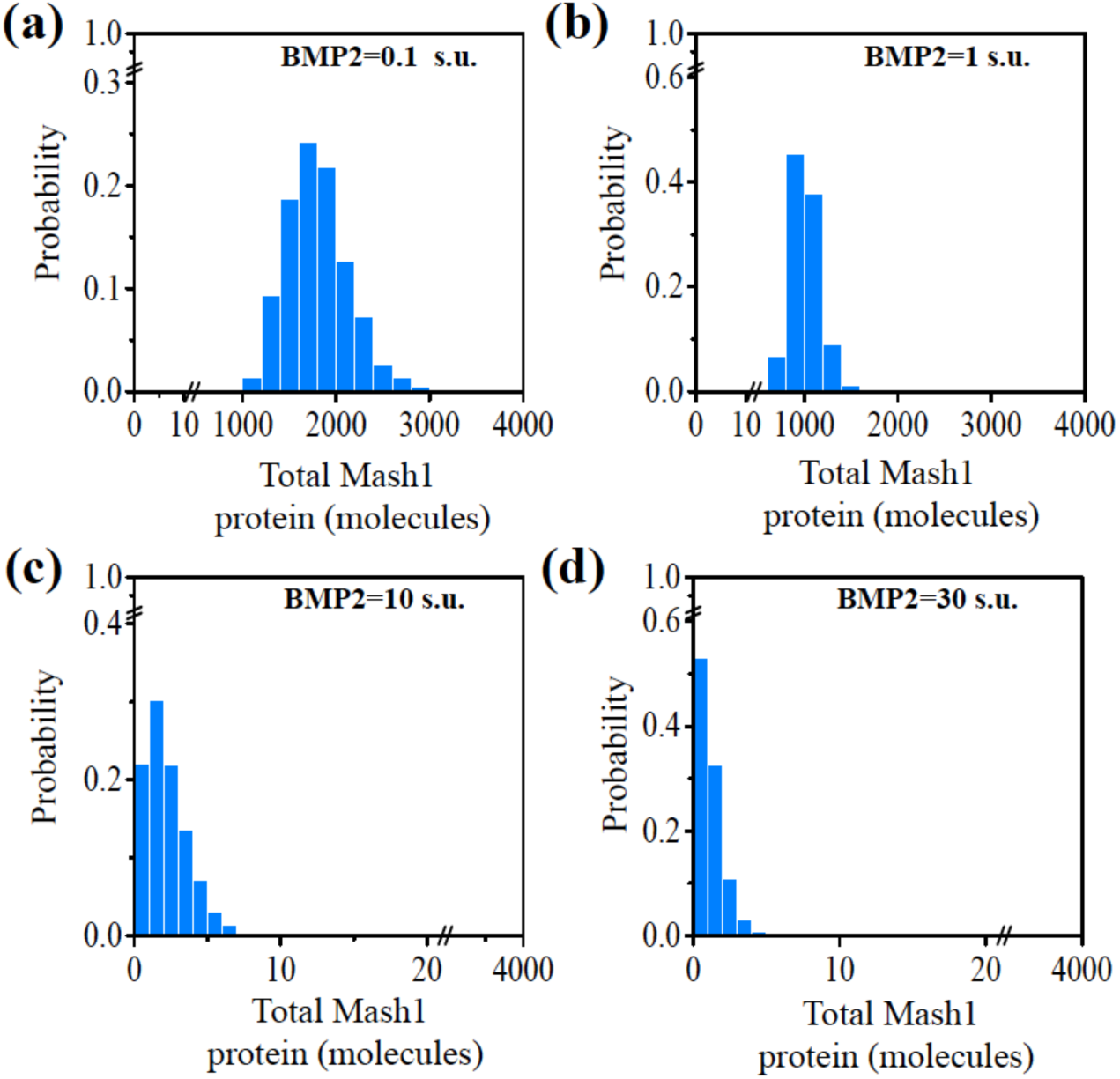
Stochastic simulation of wild type (WT) NSC's of CNS origin for 2000 cells. Steady state distributions of total Mash1 protein are plotted at different doses of BMP2 for 2000 cells (for details see method section). **(a)** BMP2=0.1X10^3^ molecules (representative of BMP2=0.1 s.u. (Fig. 2a)). **(b)** BMP2=1X10^3^ molecules (representative of BMP2=1 s.u. (Fig. 2a)). **(c)** BMP2=10X10^3^ molecules (representative of BMP2=10 s.u. (Fig. 2a)). **(d)** BMP2=30X10^3^ molecules (representative of BMP2=30 s.u. (Fig. 2a)). Developmental fate changes from neurogenic state (high Mash1 expressing cells) to gliogenic state (low Mash1 expressing cells) with increase in BMP2 dose.

**Fig. S3.**
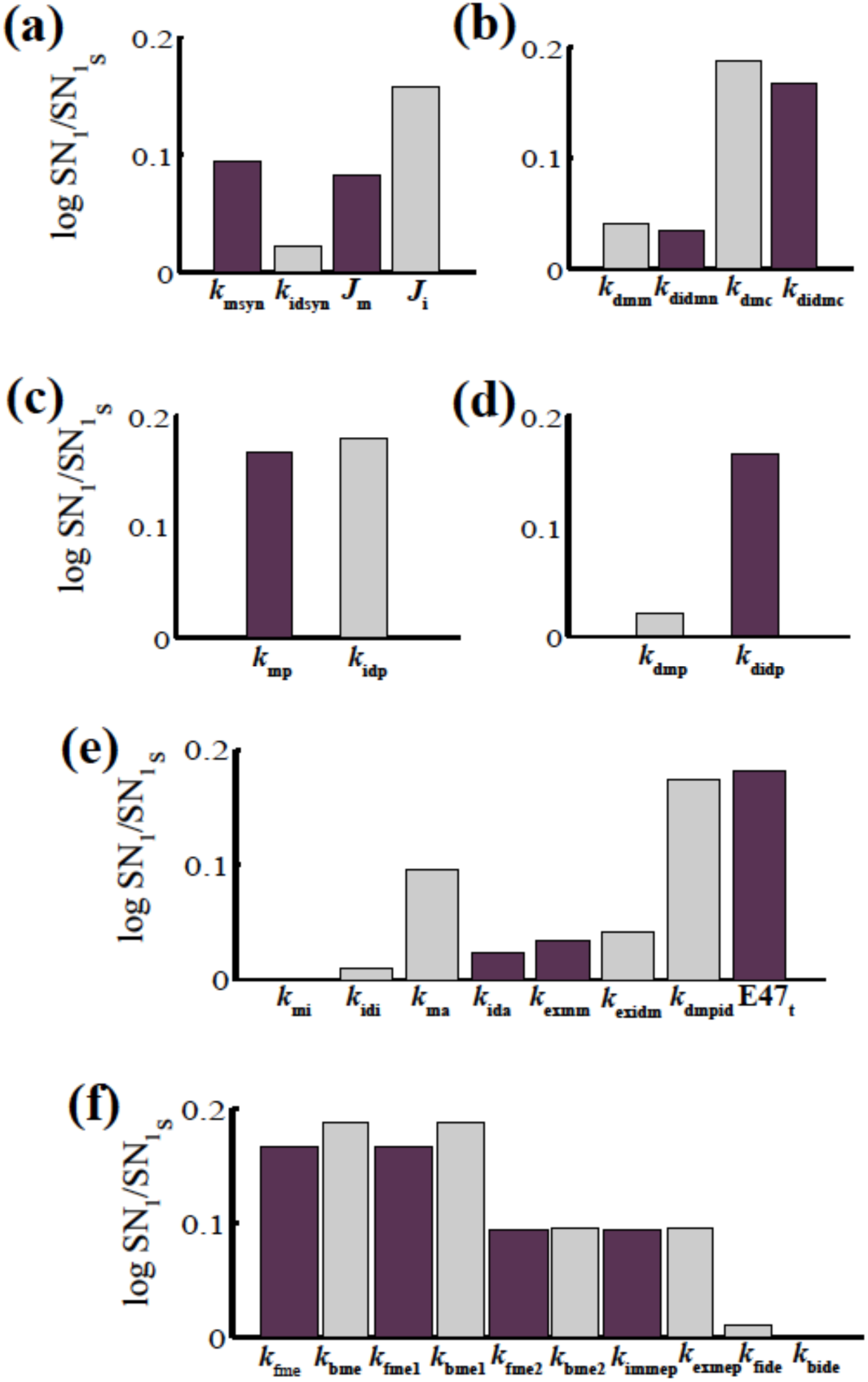
Sensitivity analysis of the model-parameters considering the position of the saddle node SN_1_ (in WT CNS) as the sensitivity criteria. Sensitivity of SN_1_ towards **(a)** Transcription rates of mRNAs. **(b)** Degradation rates of the mRNAs. **(c)** Translation rates of proteins. **(d)** Degradation rates of the Proteins. **(e-f)** Rest of the parameters. Light-grey bar signifies movement of the saddle node (SN_1_) towards higher BMP2 and dark-purple bar signifies movement of the saddle node (SN_1_) towards lower BMP2 with respect to the WT CNS situation. Sensitivity analysis was done by increasing each parameter individually by an amount of 20% of the corresponding parameter value used in the WT case (Table S2) keeping all other parameters constant.

**Fig. S4.**
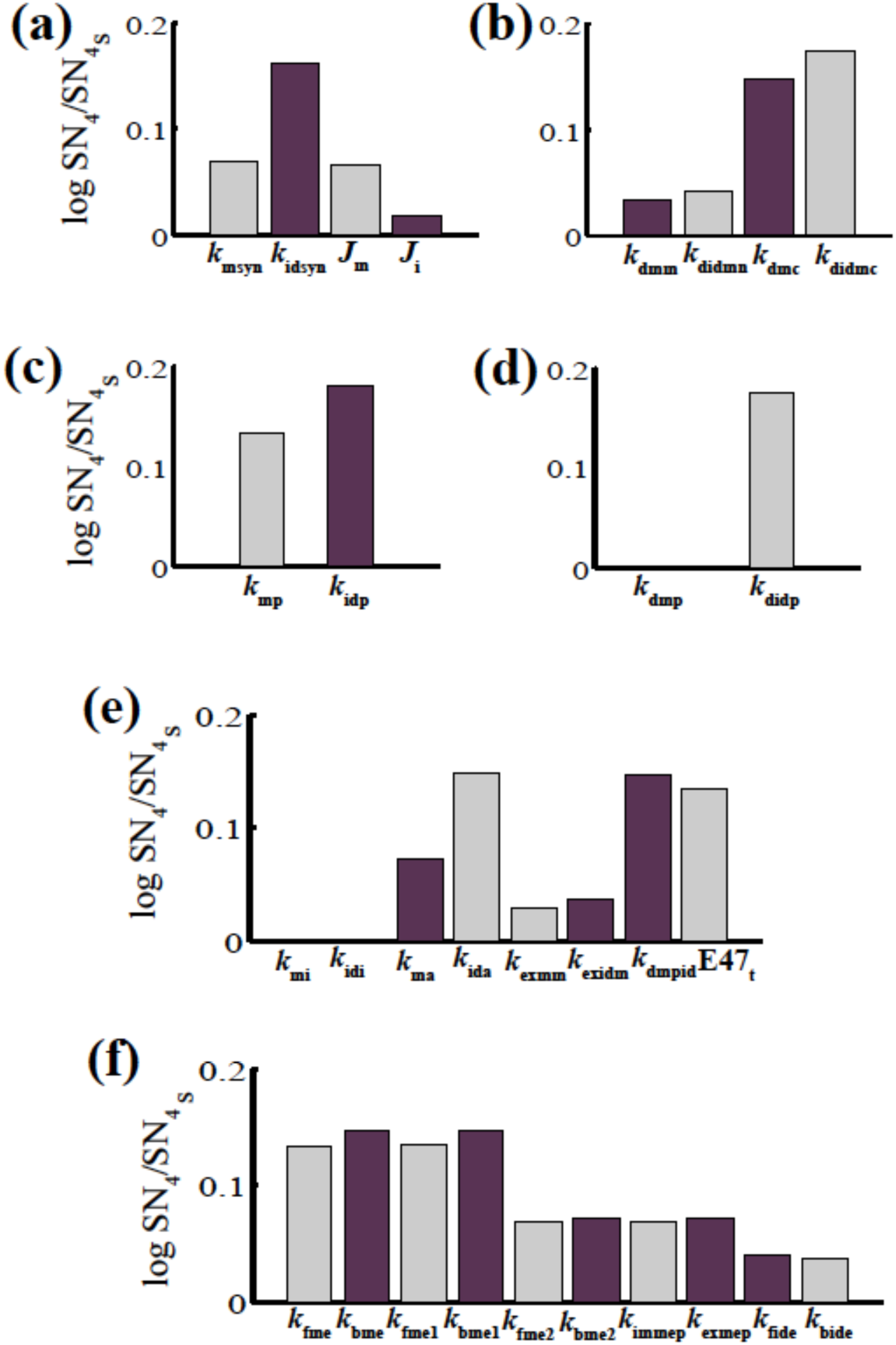
Sensitivity analysis of the model-parameters considering the position of the saddle node SN_4_ (in WT CNS) as the sensitivity criteria. Sensitivity of SN_4_ towards **(a)** Transcription rates of mRNAs. **(b)** Degradation rates of the mRNAs. **(c)** Translation rates of proteins. **(d)** Degradation rates of the Proteins. **(e-f)** Rest of the parameters. Light-grey bar signifies movement of the saddle node (SN_4_) towards higher BMP2 and dark-purple bar signifies movement of the saddle node (SN_4_) towards lower BMP2 with respect to the WT CNS situation. Sensitivity analysis was done by increasing each parameter individually by an amount of 20% of the corresponding parameter value used in the WT case (Table S2) keeping all other parameters constant.

**Fig. S5.**
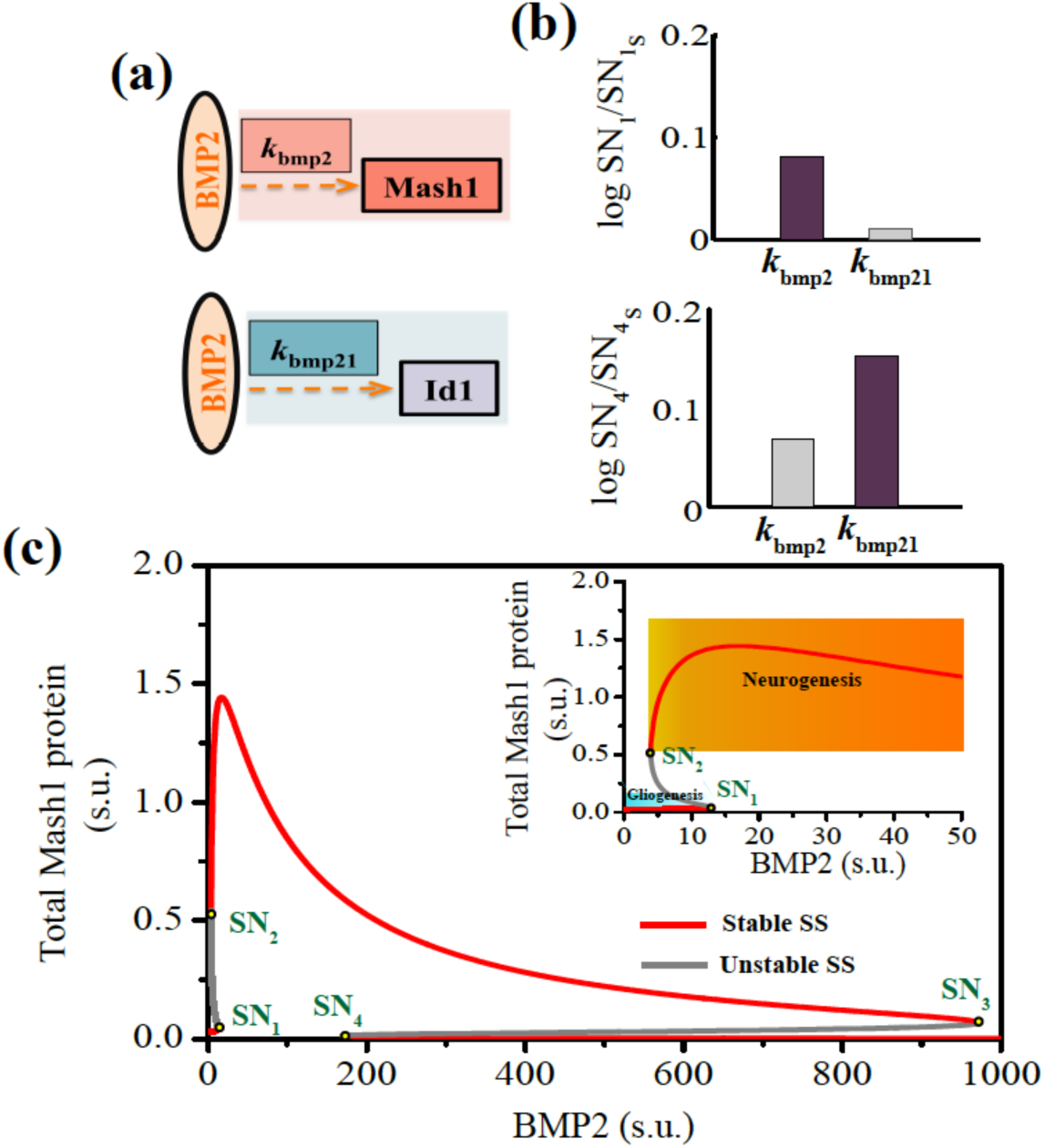
Sensitivity analysis predicts the possibility of reconciling the BMP2 driven developmental fate of WT NSC's of PNS origin. **(a)** Upper and lower panels show schematic representation of *k*_bmp2_ and *k*_bmp21_ as the transcriptional activation rates of Mash1 and Id1 by BMP2 respectively. **(b)** Upper and lower panels show sensitivities of *k*_bmp2_ and *k*_bmp21_ towards SN_1_ and SN_4_ (in Fig. 2) for WT NSC's in CNS. *k*_bmp2_ is more sensitive towards SN_1_ and *k*_bmp21_ is more sensitive towards SN_4_. Light-grey bar signifies movement of the saddle node towards higher BMP2 and dark-purple bar signifies movement of the saddle node towards lower BMP2 than the WT CNS case. Both the parameters are increased individually at an amount of 20% of the model parameters (Table S2) keeping all other parameters constant. **(c)** Bifurcation diagram of total Mash1 protein is plotted as a function of BMP2 (both the axes are shown in linear scale) in PNS. Increasing level of BMP2 switches the developmental cell fate from gliogenic state (low Mash1 expression level, shown by the graded blue region (inset)) to neurogenic state (high Mash1 expression level, shown by the graded orange region (inset)). *k*_bmp2_=400 min^−1^ and *k*_bmp21_=1.2X10^−2^ min^−1^ are used to get PNS like features. Other parameters are same as depicted in Table S2.

**Fig. S6.**
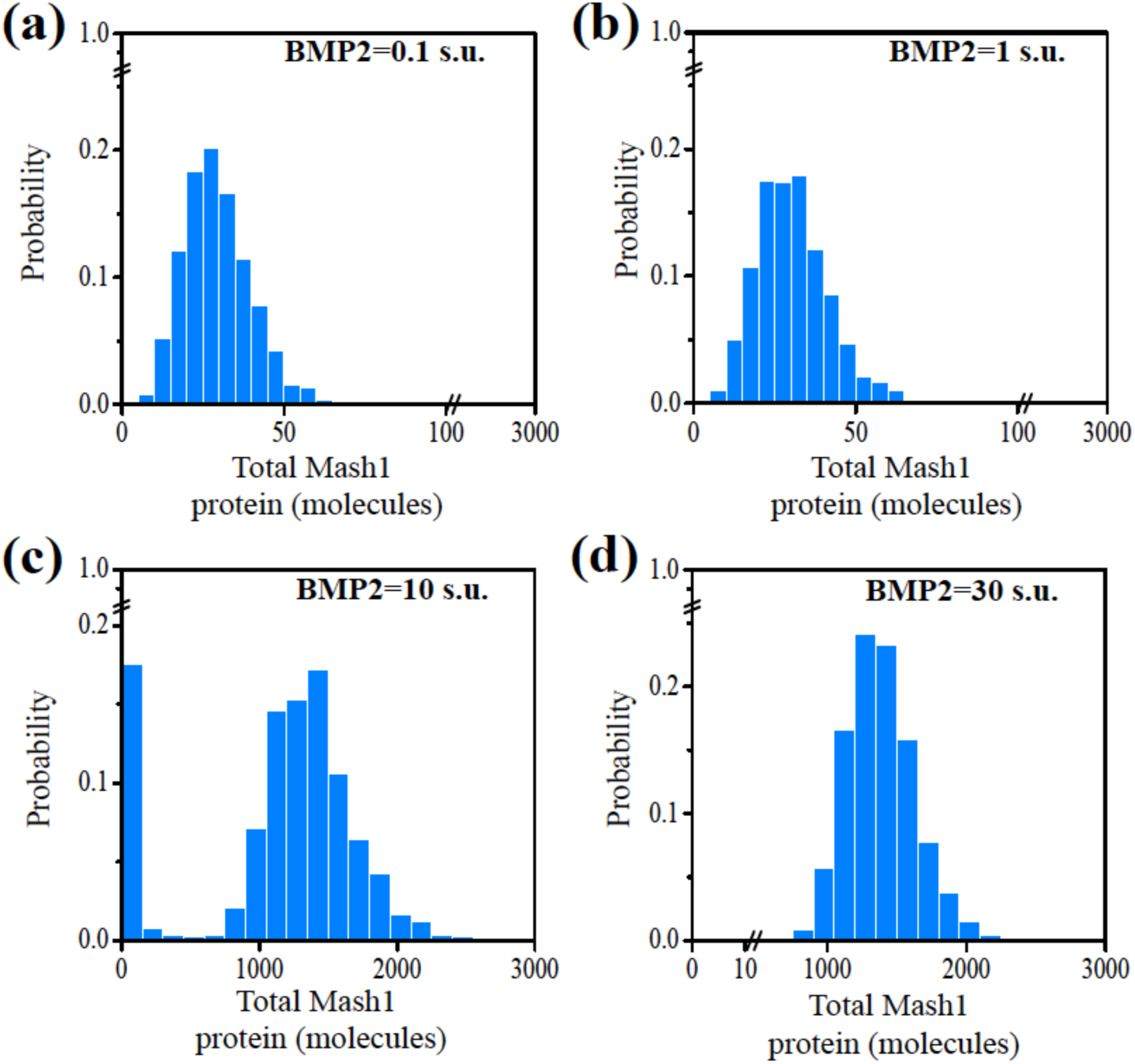
Stochastic simulation of wild type (WT) NSC's of PNS origin for 2000 cells. Steady state distributions of total Mash1 protein are plotted at different doses of BMP2 for 2000 cells (for details see method section). **(a)** BMP2=0.1X10^3^ molecules (representative of BMP2=0.1 s.u. (Fig. 3a)). **(b)** BMP2=1X10^3^ molecules (representative of BMP2=1 s.u. (Fig. 3a)). **(c)** BMP2=10X10^3^ molecules (representative of BMP2=10 s.u. (Fig. 3a)). **(d)** BMP2=30X10^3^ molecules (representative of BMP2=30 s.u. (Fig. 3a)). Developmental fate changes from gliogenic state (low Mash1 expressing cells) to neurogenic state (high Mash1 expressing cells) with increase in BMP2 dose.

**Fig. S7.**
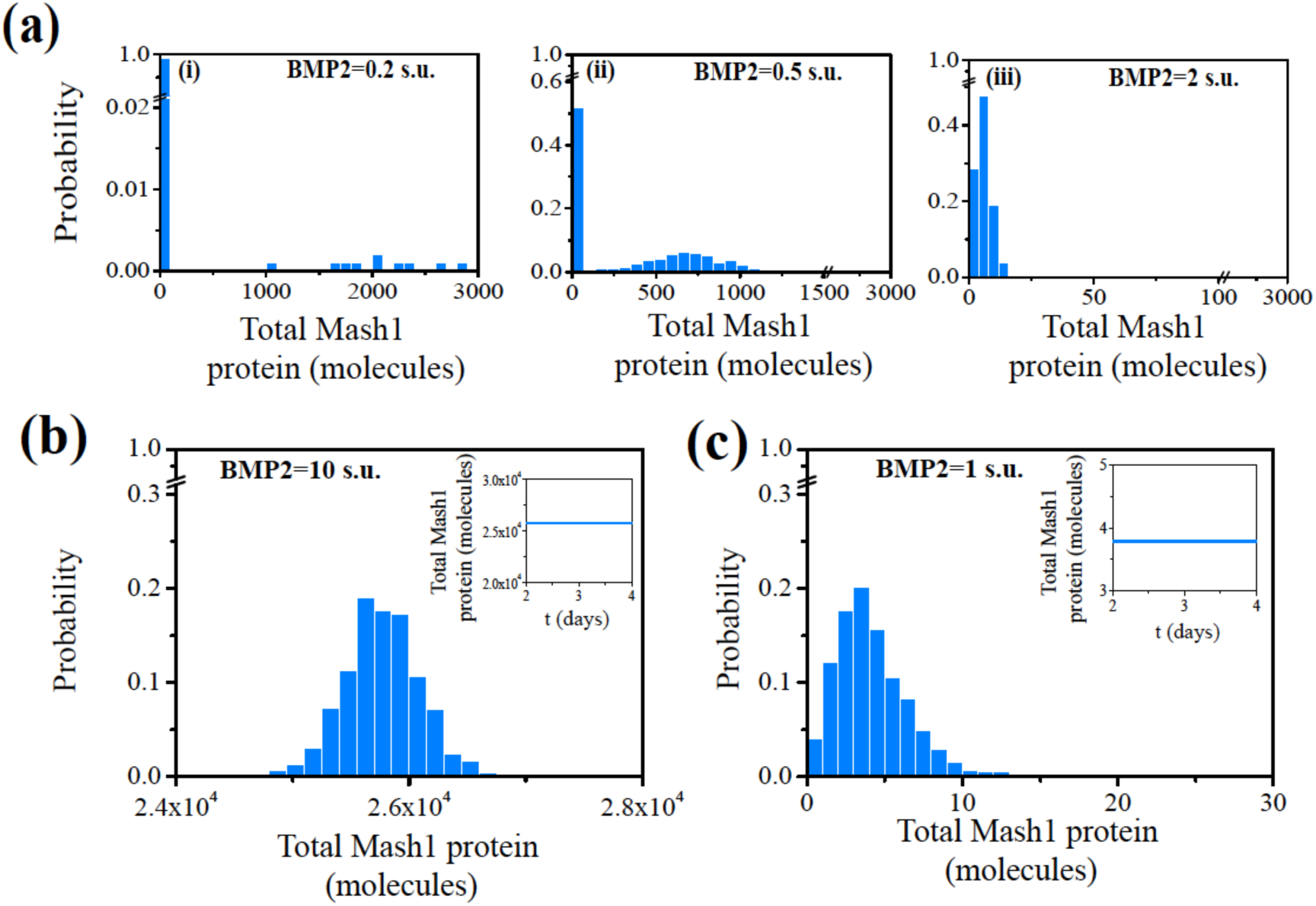
The model simulates different phenotypic conditions for NSC's in CNS. **(a)** Stochastic simulations are done at different BMP2 doses for situation 2 (Fig. 4a, E47_t_ knocked-out condition) in CNS starting from the initial condition of WT CNS of that particular BMP2 dose. **(b)** Deletion of *Id1* gene results in the up-regulation of Mash1, which indicates neurogenic fate commitment at high level of BMP2 (BMP2=10X10^3^ molecules) stochastically and deterministically (BMP2=10 s.u., inset). **(c)** Overexpression of *Id1* gene (*G*_idt_=3 X WT) results in the down-regulation of Mash1 level, which leads to gliogenesis at low dose of BMP2 stochastically (BMP2=1X10^3^ molecules) and deterministically (BMP2=1 s.u., inset). All the simulations are done for 1000 cells (for details see method section).

**Fig. S8.**
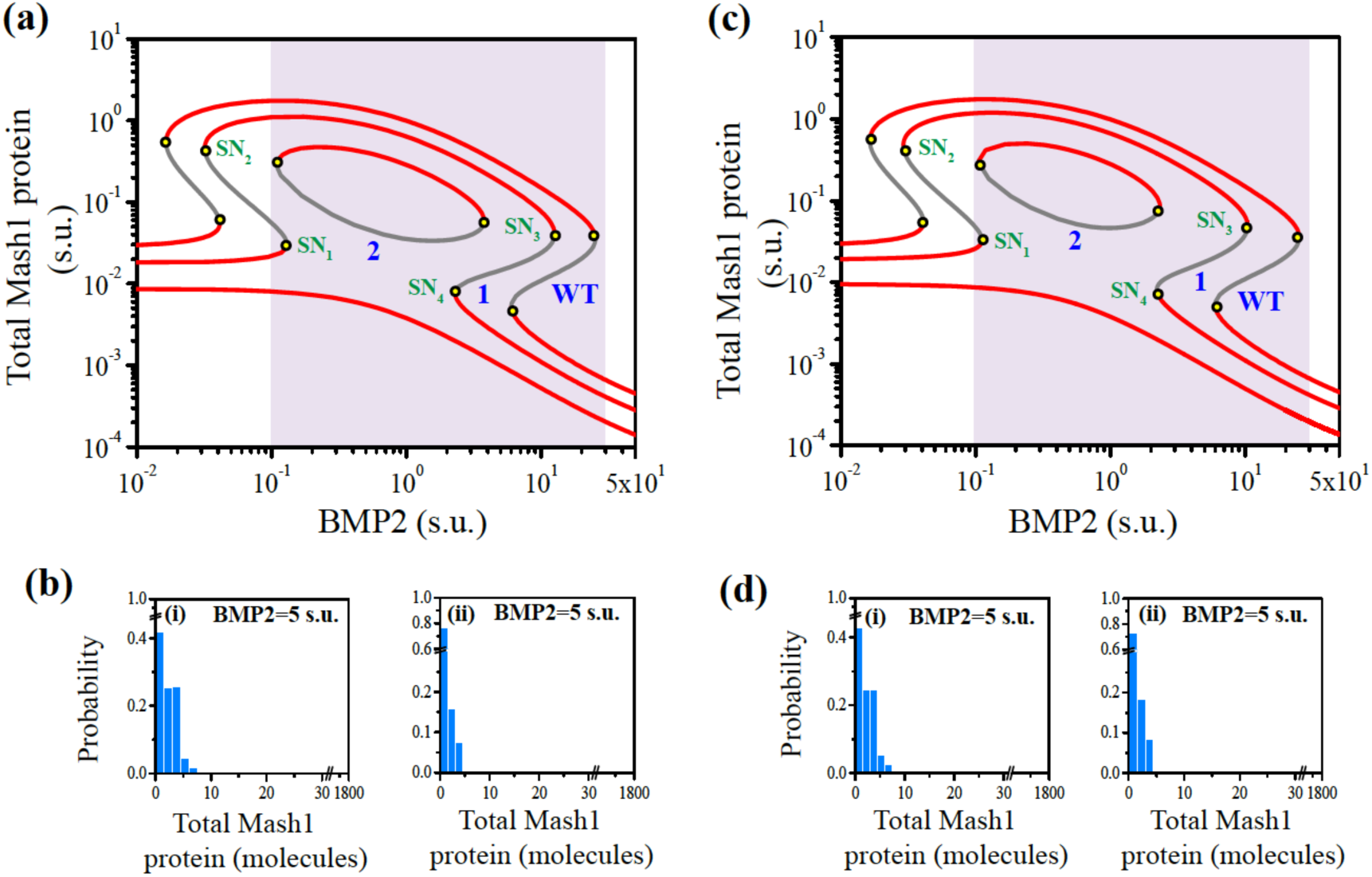
Model predicts the possible routes of unidirectional cell fate commitment of NSC's in CNS. **(a)** Decrease in *k*_mp_ leads to change in the dynamics of total Mash1 protein from mushroom bifurcation (situation 1, *k*_mp_=0.06 min^−1^) to ‘isola’ bifurcation (situation 2, *k*_m_p=0.03 min^−1^) with respect to WT CNS (*k*_mp_=0.0947 min^−1^). Other parameters are same as depicted in Table S2. **(b)** Stochastic simulations are done at a particular BMP2 dose in CNS (BMP2=5X10^3^ molecules, representative of BMP2=5 s.u.) for (i) situation 1 and (ii) situation 2. **(c)** Increase in *k*_idp_ leads to change in the dynamics of total Mash1 protein from mushroom bifurcation (situation 1, *k*_idp_=1.5 min^−1^) to ‘isola’ bifurcation (situation 2, *k_i_*_dp_=3.0 min^−1^) with respect to WT CNS (*k*_idp_=1.0 min^−1^). Other parameters are same as depicted in Table S2. **(d)** Stochastic simulations are done at a particular BMP2 dose in CNS (BMP2=5X10^3^ molecules, representative of BMP2=5 s.u.) for (i) situation 1 and (ii) situation 2. All the simulations are done for 1000 cells (for details see method section).

**Table S1.**
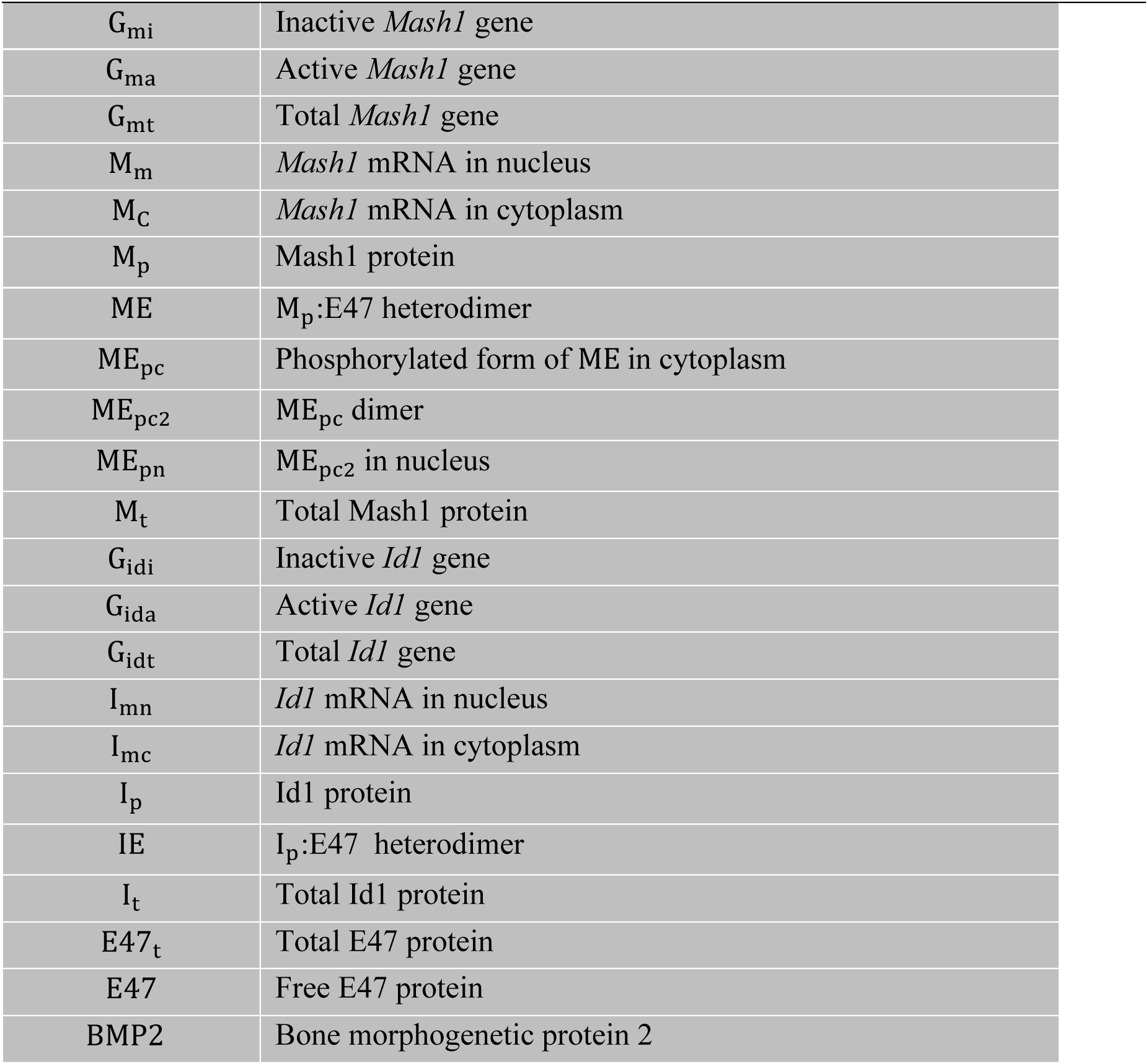
Abbreviated names of species involved in the model and their description.

**Table S2.**
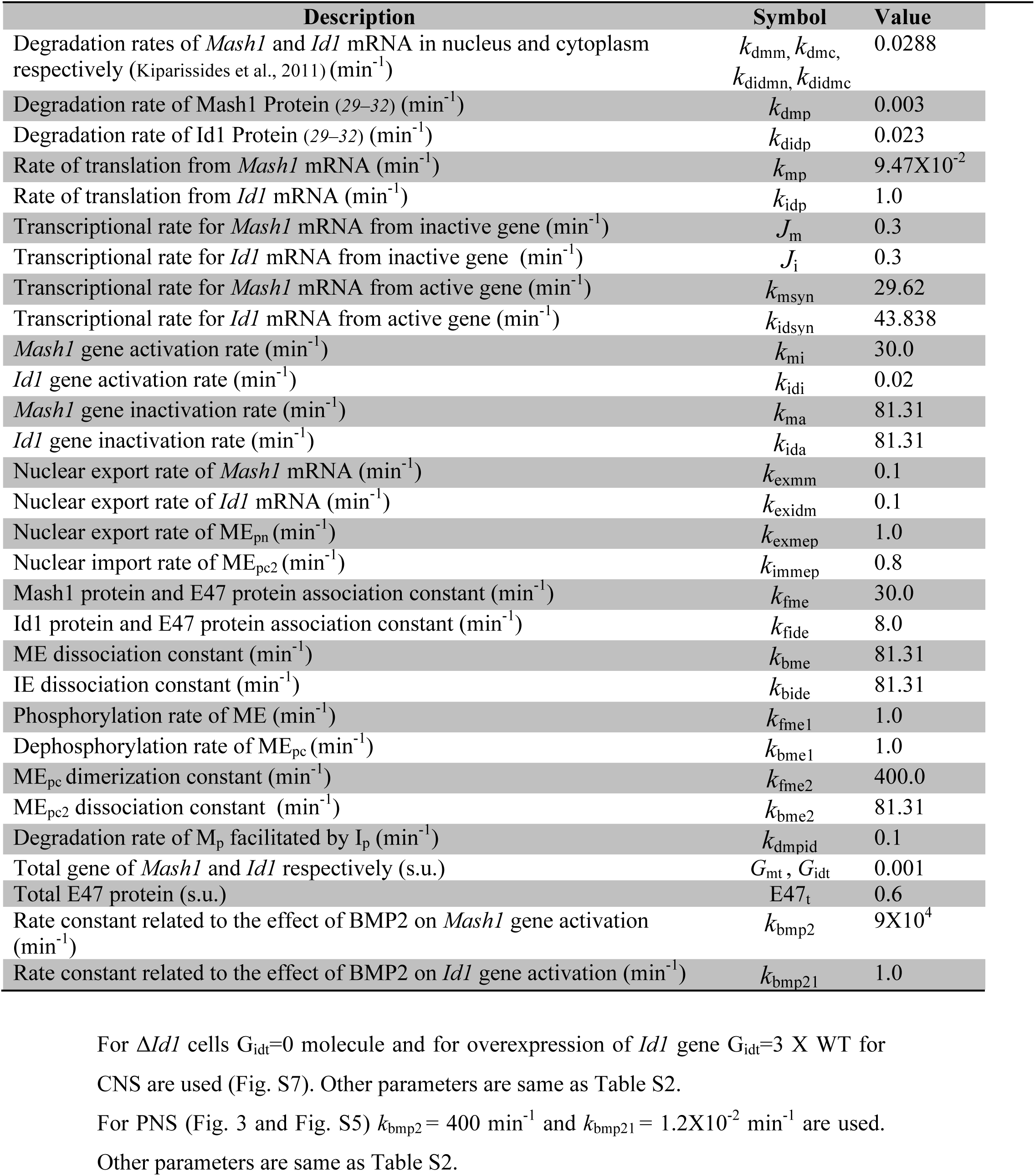
Description of the parameters, their values and sources for WT CNS.

**Table S3.** Stochastic version

| Reaction number | Reaction | Propensity of reaction |
| --- | --- | --- |
| 1 | $G_{mi} \xrightarrow{ME_{pn}} G_{ma}$ | $w_1 = k_{mi} ME_{pn} (G_{mt} - G_{ma})$ |
| 2 | $G_{mi} \xrightarrow{BMP2.ME_{pn}} G_{ma}$ | $w_2 = k_{bmp2} ME_{pn} (G_{mt} - G_{ma}) BMP2$ |
| 3 | $G_{ma} \rightarrow G_{mi}$ | $w_3 = k_{ma} G_{ma}$ |
| 4 | $\xrightarrow{G_{mi}} M_m$ | $w_4 = J_m (G_{mt} - G_{ma})$ |
| 5 | $\xrightarrow{G_{ma}} M_m$ | $w_5 = k_{msyn} G_{ma}$ |
| 6 | $M_m \rightarrow$ | $w_6 = k_{dmm} M_m$ |
| 7 | $M_m \rightarrow M_c$ | $w_7 = k_{exmm} M_m$ |
| 8 | $M_c \rightarrow$ | $w_8 = k_{dmc} M_c$ |
| 9 | $\xrightarrow{M_c} M_p$ | $w_9 = k_{mp} M_c$ |
| 10 | $M_p \rightarrow$ | $w_{10} = k_{dmp} M_p$ |
| 11 | $M_p \xrightarrow{I_p}$ | $w_{11} = k_{dmpid} I_p M_p$ |
| 12 | $M_p + E47 \rightarrow ME$ | $w_{12} = k_{fme} M_p (E47_t - ME - ME_{pc} - 2.ME_{pc2} - 2.ME_{pn} - IE)$ |
| 13 | $ME \rightarrow M_p + E47$ | $w_{13} = k_{bme} ME$ |
| 14 | $ME \rightarrow ME_{pc}$ | $w_{14} = k_{fme1} ME$ |
| 15 | $ME_{pc} \rightarrow ME$ | $w_{15} = k_{bme1} ME_{pc}$ |
| 16 | $ME_{pc} + ME_{pc} \rightarrow ME_{pc2}$ | $w_{16} = k_{fme2} ME_{pc} (ME_{pc} - 1)$ |
| 17 | $ME_{pc2} \rightarrow ME_{pc} + ME_{pc}$ | $w_{17} = k_{bme2} ME_{pc2}$ |
| 18 | $ME_{pc2} \rightarrow ME_{pn}$ | $w_{18} = k_{immep} ME_{pc2}$ |
| 19 | $ME_{pn} \rightarrow ME_{pc2}$ | $w_{19} = k_{exmep} ME_{pn}$ |
| 20 | $G_{idi} \rightarrow G_{ida}$ | $w_{20} = k_{idi} (G_{idt} - G_{ida})$ |
| 21 | $G_{idi} \xrightarrow{BMP2} G_{ida}$ | $w_{21} = k_{bmp21} (G_{idt} - G_{ida}) BMP2$ |
| 22 | $G_{ida} \rightarrow G_{idi}$ | $w_{22} = k_{ida} G_{ida}$ |
| 23 | $\xrightarrow{G_{idi}} I_{mn}$ | $w_{23} = J_i (G_{idt} - G_{ida})$ |
| 24 | $\xrightarrow{G_{ida}} I_{mn}$ | $w_{24} = k_{idsyn} G_{ida}$ |
| 25 | $I_{mn} \rightarrow$ | $w_{25} = k_{didmn} I_{mn}$ |
| 26 | $I_{mn} \rightarrow I_{mc}$ | $w_{26} = k_{exidm} I_{mn}$ |
| 27 | $I_{mc} \rightarrow$ | $w_{27} = k_{didmc} I_{mc}$ |
| 28 | $\xrightarrow{I_{mc}} I_p$ | $w_{28} = k_{idp} I_{mc}$ |
| 29 | $I_p \rightarrow$ | $w_{29} = k_{didp} I_p$ |
| 30 | $I_p + E47 \rightarrow IE$ | $w_{30} = k_{fide} I_p (E47_t - ME - ME_{pc} - 2.ME_{pc2} - 2.ME_{pn} - IE)$ |
| 31 | $IE \rightarrow I_p + E47$ | $w_{31} = k_{bide} IE$ |

By applying an appropriate conversion factor (c=1000) stochastic simulations were performed.

## SI Text

### Probable potential avenues to achieve the developmental features of WT NSC's in PNS

The model, proposed herein, performs adequately to qualitatively elucidate the developmental fate choice of NSC's of CNS under different doses of BMP2 for varied biological conditions. Now the challenge is to assess whether the same mathematical model can qualitatively portray the unique features of the WT NSC's obtained from PNS for different doses of BMP2 (i.e., low dose of BMP2 preferably leads to gliogenesis and higher dose of BMP2 leads to neuronal fate commitment) or not? Since, in this article we are more interested to understand BMP2-mediated differentiation dynamics, we concentrate on the BMP2 related parameters *k*_bmp2_ and *k*_bmp21_ in the model (Fig. S5a). As from recent theoretical advances (*21*) intuitively one can confer that the positions of the saddle nodes SN_1_ and SN_4_ critically govern the dynamics of the system as a function of BMP2 dose, in this regard we have done systematic sensitivity analysis of *k*_bmp2_ and *k*_bmp21_ by taking the position of the saddle nodes SN_1_ and SN_4_ (Fig. S5b) separately as sensitivity criteria (For details see methods section). From Fig. S5b it is evident that the saddle node SN_1_ is more sensitive with respect to the parameter *k*_bmp2_ than saddle node SN_4_, whereas the situations are just the reverse in case of the sensitivities of SN_1_ and SN_4_ as a function of *k*_bmp21_. Increase in *k*_bmp2_ leads to shift of the SN_1_ towards lower BMP2 level whereas SN_4_ moves towards higher BMP2 level. The opposite phenomenon is observed for *k*_bmp21_. Thus decrease in *k*_bmp21_ will lead to movement of SN_4_ towards higher BMP2 level to a significant extent than movement of SN_1_ towards lower BMP2 level. On the contrary, decrease in *k*_bmp2_ will result shifting of SN_1_ towards higher BMP2 level to sufficient extent than SN_4_ to lower BMP2 level. Thus decrease in *k*_bmp2_ and *k*_bmp21_ together results in the movement of both SN_1_ and SN_4_ towards higher BMP2 level, which essentially leads to PNS like behavior of the differentiation regulation under similar experimental range of BMP2 doses as observed in the case of CNS. Our model successfully predicts that by reducing both *k*_bmp2_ and *k*_bmp21_ to a required amount from the values used to delineate the WT type case of the CNS, we can have the bifurcation diagram (Fig. 3a and Fig. S5c) describing the WT situation of the NSC's obtained from PNS under different doses of BMP2. Under this situation, at low level of BMP2 the system leans towards gliogenesis (with low level of total Mash1) and higher level of BMP2 will cause neurogenesis (with higher level of total Mash1). Thus by tuning only these two parameters one can reconcile the PNS like features (rest of the parameters are same as Table S2, equations are given in Table 1).

### Biological relevance of the sensitivity analysis approach taken to reconcile PNS like behavior

The biological significance of reducing *k*_bmp2_ and *k*_bmp21_ to attain PNS like situation is quite intriguing and complex. BMP2 promotes the activation of HMG-box factor Sox10, which further induces the transient activation of *Mash1* gene (6, 37). Moreover, it had been observed experimentally that Mash1 protein negatively controls the expression level of Sox10 protein either by direct or indirect fashion(6,21,37). In our model all these regulations are acquired by the parameter *k*_bmp2_. The activity of BMP2 is mediated by its downstream transcription factor Smad, which can further regulate Id1 expression level(1, 38, 39). So changes in the expression level of Smad signaling pathway can eventually change the course of the differentiation mode of NSC's in a PNS like system, which is represented in our model in terms of *k*_bmp21_ parameter.

